# Bioenergetic responses to β-adrenergic stimulation in beige adipocyte depend on actomyosin driven forces

**DOI:** 10.1101/2025.11.26.690550

**Authors:** Yuchen He, Lu Ling, Garrett Dempsey, Nayiri Kalindjian, Zoie Verghese, Erzhen Chen, Irene Liparulo, Isabel Cho, Vyvylyn Tran, Sanjay Kumar, Andreas Stahl

## Abstract

Beige (BeAT) and white adipoctyes (WAT) reside within the same fat depots but differ in their responses to β-adrenergic stimulation. While both activate classical PKA-mediated pathways, only beige adipocytes show induction of UCP1 expression and enhanced mitochondrial respiration, suggesting additional mechanisms that distinguish their responses. Here we uncover a parallel, non-canonical biomechanical pathway specific to beige adipocytes that is essential for thermogenic activation. Mechanistically, β-AR stimulation induces a rapid Myh9-mediated actomyosin contraction and stiffening in beige, but not white, adipocytes that activates focal adhesion kinase (FAK), which we show to be critical for the expression of oxidative and thermogenic genes including UCP1 and PLIN5. These findings reveal a new biomechanical Myh9-FAK signaling arm downstream of b-adrenergic activation that differentiates thermogenic from white adipocytes.

## Introduction

Obesity is a rapidly growing global epidemic^1^ and major risk factor for various metabolic diseases, including type-2 diabetes mellitus^2^, nonalcoholic fatty liver disease^3^, and cardiovascular diseases^4^. A better understanding of adipose tissue homeostasis, particularly microenvironmental signals that regulate the balance between storage vs consumption of lipids, may provide mechanistic and, potentially, therapeutic insights for combatting obesity and associated co-morbidities^5^. Subcutaneous adipose tissues contain white and beige adipocytes. While white adipocytes primarily serve lipid storage, brown and beige adipocytes are thermogenic and characterized by high mitochondrial content and multilocular lipid droplets, enabling them to dissipate energy as heat via uncoupled mitochondrial respiration facilitated by uncoupling protein 1 (UCP1)^6^. Comparing to classical brown adipocytes, which are innate and diminish with age, beige adipocytes are dispersed within white adipose tissues through differentiation and trans-differentiation of progenitor cells and white adipocytes^7–9^ and are thought to represent the majority of thermogenic tissue in adult humans^10^.

Several stimuli can activate BeAT with b-adrenergic signaling representing the most important physiological trigger^6,9^. During cold exposure, sympathetic nerves release norepinephrine into adipose depots, rapidly activating β-AR signaling through the Gαs - .adenylyl cyclase - cAMP - protein kinase A (PKA) cascade^11^. In white adipose tissue, activated PKA phosphorylates hormone-sensitive lipase (HSL)^12,13^, mobilizing non-esterified fatty acids and glycerol to support peripheral energy demands^14,15^. In contrast, BeAT and BAT acutely increase fatty-acid uptake and oxidation while engaging UCP1-mediated mitochondrial uncoupling to generate heat^6^ illustrated by the finding that genetic disruption of FATP1 (Slc27a1) impairs free fatty acids uptake and non-shivering thermogenesis in brown adipose tissue^16^. Additionally, PKA phosphorylates cAMP responsive element binding protein (CREB), which enhances UCP1 transcription in BeAT and BAT^17^.

We previously showed a role for biomechanical signaling in classical brown adipose tissue that depends on mysosin^18^. Myosins are classified both in terms of gene name for their heavy-chain subunits (e.g. Myh9) and functional behavior (e.g. Type II)^19^. Type II myosins are hexameric complexes^18^ composed of two myosin heavy chains, two myosin regulatory light chains, and two myosin essential light chains^20^. Traditionally studied in the context of cell structure, mechanics, and motility, myosin II-based forced sensing and force generation are increasingly implicated in the regulation of an expanse of cellular activities such as metabolism^21^ and stem cell fate commitment^22^. Other studies also suggest that actomyosin network reorganization can also directly influence mitochondrial dynamics^23–25^, underscoring the potential of subcellular actomyosin networks in regulating key functions involved in metabolic homeostasis^26–28^. In BAT Myh7 mediates a YAP/TAZ-dependent signaling pathway to enhance thermogenic capacity and to increase metabolic rates^18^. However, Myh7 is not expressed by BeAT^8,18,29^, potentially due to developmental differences between classical BAT and WAT interspaced BeAT. Overall, it remains unknown whether BeAT metabolism is regulated by biomechanical signaling pathways and whether alternate myosin isoforms underlie that regulation.

In this study, we identify a non-canonical, actomyosin-mediated pathway specific to BeAT involving β-adrenergic–induced cytosolic Ca²⁺ elevation, Myh9-dependent tension generation, and focal adhesion kinase (FAK) activation. This mechanical signaling axis regulates beige adipocyte UCP1 expression and LD-mitochondria mediated bioenergetics, in part through the expression of the LD-associated protein PLIN5. Disruption of this pathway impairs thermogenesis and lipid metabolism, highlighting a novel mechanism linking cytoskeletal tension to metabolic function in beige adipocytes.

## Results

### Beige adipocyte activation is coupled with increased actomyosin-mediated tension

To assess WAT mechanical properties during thermogenesis induction, we used atomic force microscopy (AFM) to measure the Young’s modulus of inguinal white adipose tissue (iWAT) from mice housed at room temperature (23°C) or subjected to cold exposure (4°C). Cold exposure for 7 days significantly increased the mean iWAT tissue stiffness from ∼700 Pa to ∼2.7 kPa (Fig 1a, b), suggesting enhanced mechanical tension during thermogenic activation.

**Figure 1.**
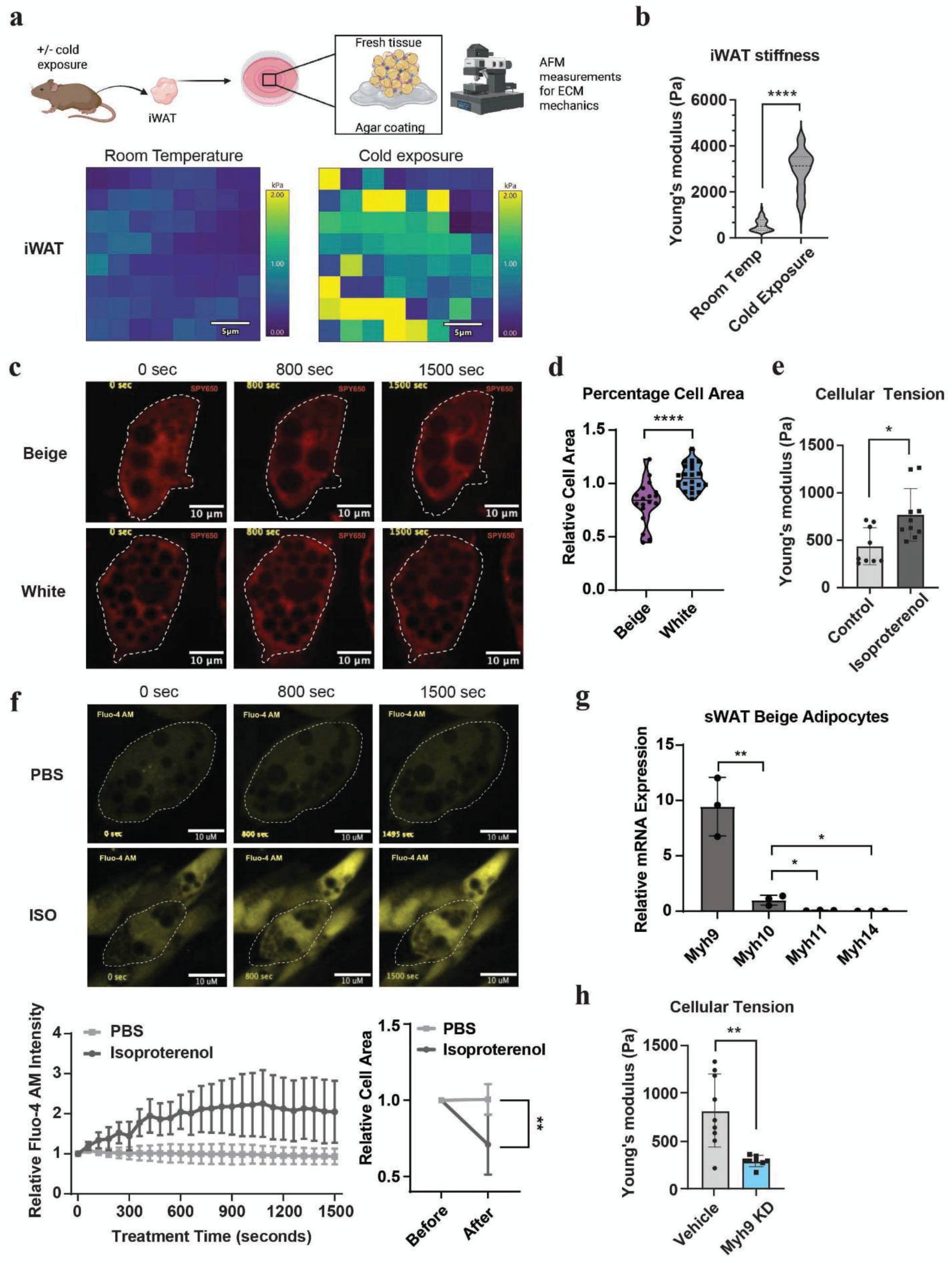
BeAT activation is associated with increased actomyosin-mediated intracellular tension. (a) Schematic showing sample harvesting process for AFM measurements of isolated inguinal white adipose tissue (IWAT) of mice housed at either room temperature or 4°C (cold exposure). Heat map showing stiffness distribution within tissue explants at room temperature vs 4°C (b) AFM measurements of tension (Young’s modulus) of tissues at room temperature vs 4°C. n =4 biological samples from each group. (c) Cellular contraction of SVF-derived beige and white adipocytes after 1 μM isoproterenol treatment. (d) Cell area change quantification of d, relative cell area is defined by the ratio of cell area at the end of experiment to cell area at the beginning of the experiment (n=18 or 22, 2 independent batches of experiments). (e) AFM measurements of cellular stiffness of in vitro beige adipocytes 2 hr after isoproterenol simulation. (f) Fluorescent imaging of cytosolic Ca2+ of sWAT beige adipocytes measured with 4 μM of Fluo-4 AM after 1 μM isoproterenol treatment. Below shows mean fluorescent intensity of Fluo 4 AM indicated cytosolic Ca2+ and relative cell area changes of sWAT beige adipocytes without or with 1 μM isoproterenol treatment (n=7 or 23, 2 independent experiments). (g) Relative expression of myosin heavy chains (Myh) in sWAT beige adipocytes as measured via qPCR. (h) AFM measurements of cellular stiffness for Myh9 shRNA KD cells vs vehicle control. Error bars represent mean ± SD. ∗: p < 0.05, ∗∗: p < 0.01. ∗∗∗: p < 0.001, ∗∗∗∗: p < 0.0001

Given that both beige and white adipocytes are present in iWAT, we next sought to determine whether the observed increase in mechanical tension was specific to beige adipocytes. Re-analysis of a published single nucleus RNA-seq dataset from cold-exposed mouse iWAT revealed significantly lower expression of actomyosin-related genes in white compared to beige adipocytes^30^, suggesting that white adipocytes are less capable of actomyosin-mediated contraction than beige adipocytes (Supplementary Fig. 1). To assess cell type specificity, we differentiated stromal vascular fraction (SVF) cells under beige- or white-inducing adipogenic conditions and tracked morphological changes following treatment with the pan-b-AR agonist isoproterenol^31^. Interestingly, while isoproterenol induced an average reduction in cell area of 18% in beige adipocytes, no change in cell sizes were observed in white adipocytes (Fig. 1c, d), indicating that β-adrenergic–induced cellular shape changes were specific to beige adipocytes. We further validated this finding using an immortalized *in vitro* beige adipocyte line (sWAT), where isoproterenol treatment markedly increased mature beige adipocyte stiffness from ∼400 Pa to ∼800 Pa (Fig. 1e), mirroring the tissue-level stiffening observed *in vivo*. Elevated cytosolic calcium concentration, which correlates with actomyosin contractility, has been previously linked with b-adrenergic signaling in beige adipocytes^32^. Consistent with previous reports, isoproterenol treatment significantly increased beige adipocyte intracellular calcium (Fig 1f). Associated with cytosolic calcium increase, a reduction in cell size was also observed in calcium imaging experiments.

Given the relevance of calcium and myosin in contractile activity, we examined the expression of myosin heavy chain isoforms in sWAT beige adipocytes (Fig 1g) and iWAT tissue (Supplementary Fig 2a). Myh9, which encodes the heavy chain of non-muscle myosin IIA (NM-IIA), emerged as the most abundant myosin isoform in both analyses. Myh9 binds to actin filaments to generate forces and is involved in a variety of cell types and processes such as maintenance of cell shape, adhesion, and migration^33,34^. shRNA knockdown (KD) of Myh9 significantly decreased cellular stiffness in beige adipocytes (Fig. 1h), confirming its role in force generation in this context. Myh10, which encodes the heavy chain of non-muscle myosin IIB (NMIIB)^35^, was the second most abundant isoform in the screen, expressed at less than one third of the level of Myh9. However, shRNA KD of Myh10 had no significant impact on cellular stiffness (Supplementary Fig 2b, c), and no significant changes in UCP1 expression and cellular respiration were observed in Myh10 KD cells (Supplementary Fig 2d, e). Consistent with results in the shRNA knockdown cell-lines, siRNA knockdown of Myh9 and Myh10 in SVF-derived beige adipocytes also suggested only Myh9 but not Myh10 KD caused significant decrease in UCP1 expression (Supplementary Fig 2f), motivating us to focus on Myh9 in subsequent studies.

### b-adrenergic stimulation is associated with local actomyosin remodeling around lipid droplets

In addition to whole-cell contraction, b-adrenergic stimulation also triggered localized actomyosin remodeling surrounding lipid droplets (LDs) (Fig 2a). Live imaging showed the formation of actin “cages” surrounding LDs within minutes of isoproterenol treatment (Fig 2b). Super-resolution structured illumination microscopy (SIM) further revealed a distinct Myh9–actin network encasing individual LDs (Fig 2c), hinting at a role for localized actomyosin Myh9–actin–dependent force generation in regulating LDs.

**Figure 2.**
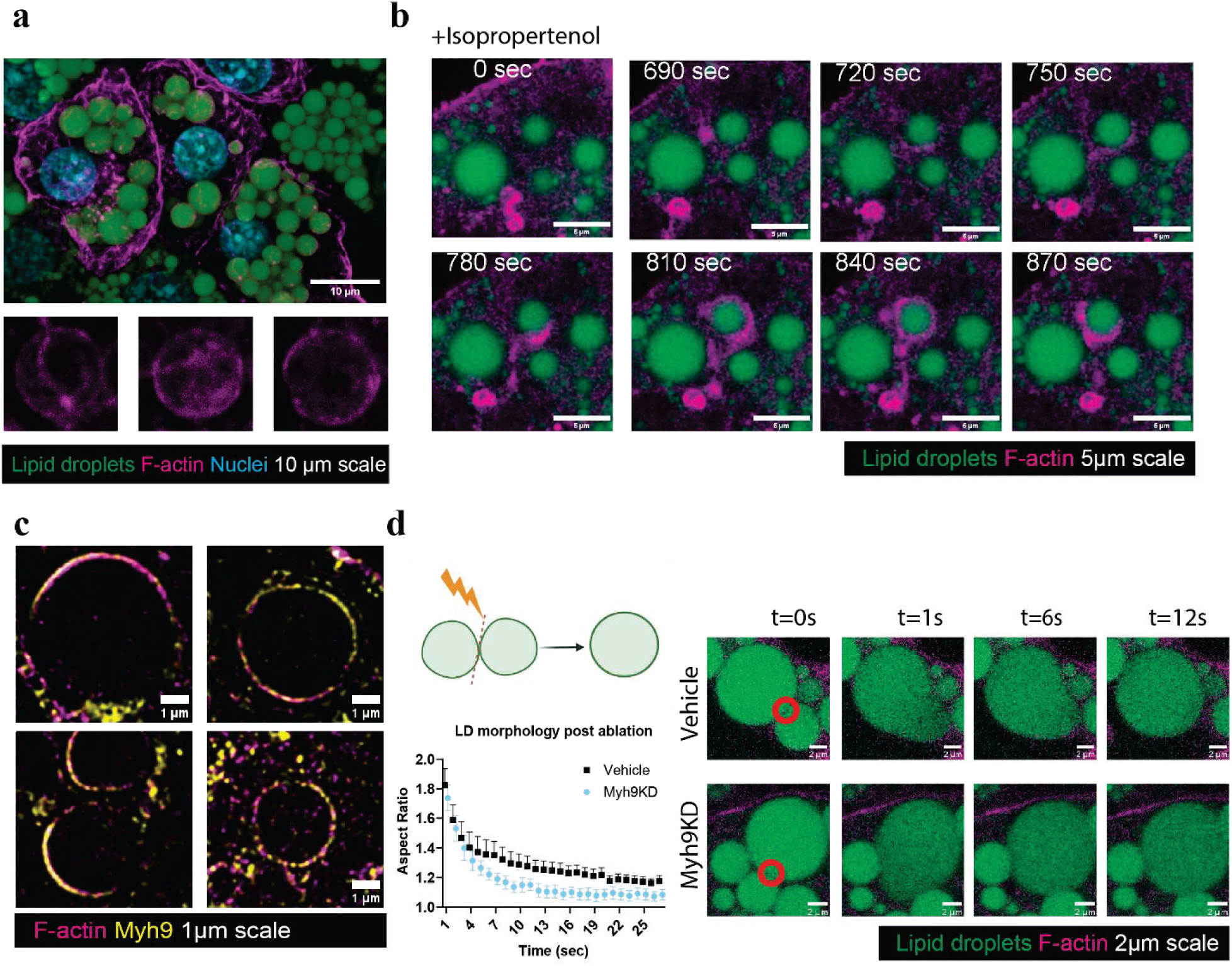
Beige adipocyte activation is coupled with actin rearrangements around lipid droplet and loss of Myh9 impairs subcellular organelle architecture and function. (a) Maximal projected super resolution image showing example of actin (magenta) around lipid droplets LD (green). Three small panel images showing the F-actin structures around a single LD. 5μm scale. (b) Time lapse images showing an example of dynamic formation of actin (magenta) around LD (green) upon isoproterenol stimulation starting from time 0 sec to 870 sec. (c) Representative SIM image showing actin (magenta) and Myh9 (yellow) distributions around lipid droplets (LD, hollow region in the center). 1 μm scale. (d) Schematic and time lapse images showing laser ablation of actin between two adjacent LDs (green) and resulting fusion events in Myh9 KD vs Vehicle cells transfected with iRFP670 Lifeact (magenta). Quantification of fusion kinetics were plotted below, n=7 in each condition. 2μm scale. Error bars represent mean ± SEM.

To investigate the local mechanical contribution of the actin network to lipid droplets (LDs), we applied laser nanosurgery to selectively sever the actin-based structures surrounding LDs^36^. Ablation of actin fibers between adjacent LDs caused these LDs to fuse within seconds, a timescale drastically faster than spontaneous fusion events, which typically occur over minutes to hours^37^. Myh9 KD cells showed accelerated LD fusion kinetics following ablation compared to controls (Fig 2d), suggesting that the Myh9–actin network localized to LDs contributes to LD integrity and stability.

### Loss of Myh9 impairs mitochondrial functions of beige adipocytes

Given that Myh9 KD resulted in reduced cellular stiffness and destabilized LDs, we next sought to evaluate its broader impact on beige adipocyte function. RNA-seq analysis of Myh9 KD beige adipocytes versus vehicle control cells revealed widespread transcriptional alterations across genes that function in multiple organelles (Fig. 3a). Among the significantly down-regulated genes, uncoupling protein 1 (UCP1) and perilipin 5 (PLIN5) stood out as key players highly associated with beige adipocyte metabolism (Fig. 3b). UCP1 directly mediates non-shivering thermogenesis, and PLIN5, a lipid droplet-associated protein,^38,39^ is one of the few known proteins that mediate LD–mitochondria contacts and facilitate fatty acid trafficking for b-oxidation^40–42^.

**Figure 3.**
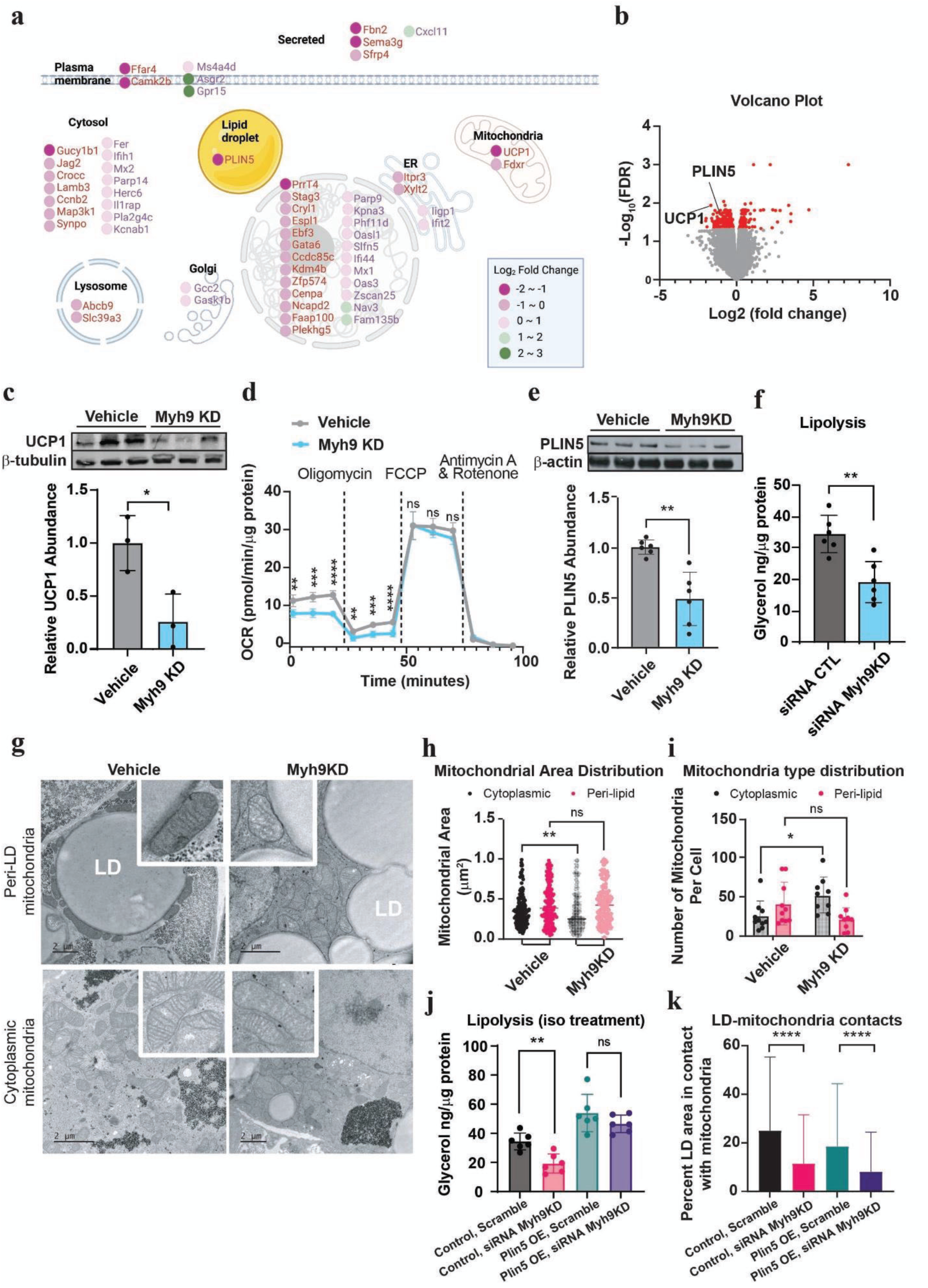
Myh9 is the predominant myosin heavy chain in regulation of beige adipocyte function. (a) Cell map showing the top 50 upregulated (purple) and 50 downregulated genes (plum) from RNA seq analysis between Myh9 KD and veh cells. (b) Volcano plot obtained from DESeq2 analysis of My9KD vs veh beige adipocytes from RNA seq analysis. n= 3 per group. (c) UCP1 protein expression assessed by western blot in inducible Myh9 KD sWAT beige adipocytes without (vehicle) or with induction of Myh9 shRNA expression for 7 days during differentiation (n=3 per group). (d) Cellular respiration of inducible Myh9 KD sWAT beige adipocytes as treated in c (n=5 per group). (e) PLIN5 protein expression assessed by western blot in inducible Myh9 KD sWAT beige. (f) Quantification of lipolysis in Myh9 shKD cells measured by free glycerol assay. n=6 samples. (g) TEM images showing mitochondria subpopulation (peri-lipid and cytosolic) and lipid droplets in vehicle control and myh9 KD cells. (h) Quantification of peri-lipid and cytosolic mitochondria sizes (0 to 1 μm^2^) in vehicle control and myh9 KD cells. (i) Quantification of peri-lipid and cytosolic mitochondria numbers per cell in vehicle control and myh9 KD cells. A minimum of 140 mitochondria per group of 10 cells were analyzed. (j) Lipolysis assay showing glycerol release from PLIN5 OE vs control cells with siRNA negative control or Myh9 siRNA KD treatment. n=6 per group. (k) Quantification of LD-mitochondria contact site area in PLIN5 OE vs control cells with siRNA negative control or Myh9 siRNA KD treatment. A minimum of 2500 LDs per group were analyzed. Error bars represent mean ± SD. ∗p < 0.05, ∗∗p < 0.01. ∗∗∗p < 0.001, ∗∗∗∗ p < 0.0001

Reductions of both UCP1 and PLIN5 expression were confirmed with western blotting (Fig 3c and e). These were accompanied by a significant reduction in cellular basal and uncoupled respiration as measured via cellular respirometry (Fig. 3d), as well as with decreased lipolysis rate in Myh9 KD cells (Fig 3f).

Mitochondria in contact with LDs (peri-LD mitochondria) have been reported in brown and beige adipocytes, where tethering of mitochondria to LDs enhances fatty acid trafficking and provides fuel for thermogenesis via b-oxidation^43,44^. To test whether reduced PLIN5 expression in Myh9KD cells is associated with altered LD–mitochondria contact sites, we used transmission electron microscopy (TEM) to visualize organelle structures (Fig 3g). TEM quantification revealed fewer peri-LD mitochondria and more cytoplasmic mitochondria in Myh9KD cells. (Fig 3h, i).

To further investigate the functional relationship between Myh9 and PLIN5, we generated PLIN5 overexpression (OE) beige adipocytes. PLIN5 OE successfully rescued the dysregulated lipolysis in beige adipocytes with Myh9 siRNA KD (Fig 3j). In line with the rescue effect of lipolysis, the maximal respiratory capacity, as reflected by cell’s ability to transport and oxidize available substrates, was improved in PLIN5 OE cells treated with Myh9 siRNA KD (Supplementary Fig. 3a). However, the decreased basal and uncoupled respiration caused by Myh9 siRNA KD was still evident in PLIN5 OE cells. Moreover, PLIN5 OE alone was not sufficient to rescue UCP1 expression in Myh9 KD cells (Supplementary Fig. 3c), suggesting the involvement of additional signaling cascades that mediate the mechanical regulation of thermogenesis programing.

Interestingly, LD–mitochondria contacts were not restored in PLIN5 OE cells treated with Myh9 siRNA KD (Fig. 3k), indicating that additional LD-mitochondria tethering mechanisms are impaired and PLIN5 alone is insufficient to restore these interactions. One possible explanation is that Myh9 is directly involved in LD–mitochondria tethering, as our immunofluorescence staining showed that Myh9 localized not only to LDs but also to mitochondria in contact with LDs (Supplementary Fig. 3d). To further explore connections between Myh9 and mitochondria organization, we performed targeted western blots to determine if Myh9 suppression altered expression or phosphorylation of specific targets such as fusion proteins Optic Atrophy 1 (OPA1) and Mitofusin 2 (MFN2)^45^ as well as fission protein dynamin-related protein 1 (Drp1)^24,25^. However, we observed no significant changes in the expression of these proteins (Supplemental Fig 3e-g). Therefore, the subcellular spatial localization of Myh9 is important for regulating LD–mitochondria structural interactions and associated lipolytic processes that are independent of canonical mitochondrial fission and fusion activities.

### Actomyosin regulation of mitochondrial gene expression is PKA-independent

In addition to above gene expression changes at the unstimulated baseline state, we observed that the Myh9 KD cells failed to induce UCP1 expression in response to b-AR stimulation. Canonically, expression of UCP1 in beige adipocytes can be induced through the PKA-CREB pathway following b-adrenergic stimulation. When treated with isoproterenol, UCP1 mRNA levels were markedly increased in control cells but increased to a much lesser extent in Myh9 KD cells (Supp Fig 4a), indicating impaired signaling downstream of b-AR. This result was supported by the finding of reduced b-AR-induced UCP1 protein in Myh9 KD cells (Supp Fig 4b), as well as decreased cellular basal and uncoupled respiration (Supp Fig 4c).

In our Myh9 KD model, Myh9 is suppressed throughout the entire course of beige adipocyte differentiation and may therefore exert long-term secondary effects. To assess the acute role of actomyosin-mediated tension in b-adrenergic signaling, we employed a pharmacological perturbation approach, inhibiting actomyosin activity with blebbistatin, which is a pan type II myosin inhibitor^46^, in mature beige adipocytes. Blebbistatin treatment effectively reduced cellular stiffness (Fig 4a) and strongly suppressed UCP1 under both unstimulated and isoproterenol-stimulated conditions (Fig 4b, c). Consistent with these transcriptional changes, blebbistatin-treated cells displayed reduced basal and uncoupled respiration under both unstimulated and isoproterenol-stimulated conditions (Fig 4d), phenocopying the defects observed in Myh9 KD cells. Conversely, pharmacological activation of actomyosin using the myosin activator EMD-57003^47^ amplified isoproterenol-induced UCP1 expression (Supplementary Fig 5), reinforcing the role of actomyosin contraction in b-adrenergic agonism regulated beige adipocyte thermogenic gene expression.

**Figure 4.**
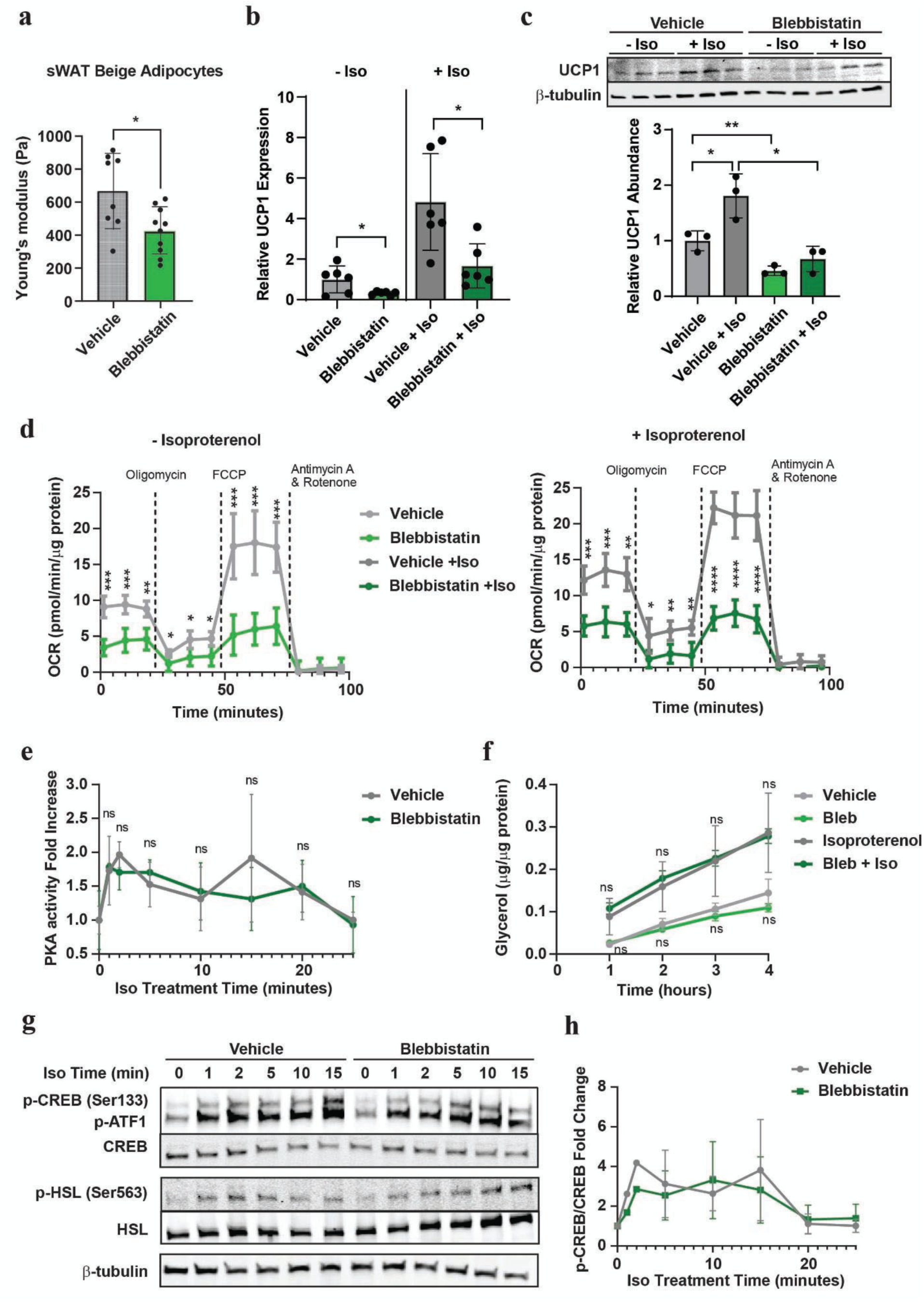
Actmyosin-mediated thermogenic gene expression regulation is independent of classical PKA mediated pathway. (a) AFM measurements of cellular stiffness for sWAT beige adipocytes treated without or with 50 mM Blebbistatin for 2 hours. (b) UCP1 mRNA expression measured by qPCR in sWAT beige adipocytes treated without (vehicle) and 1 mM isoproterenol overnight (n=3 per group). (n=6 per group). (c) UCP1 protein expression assessed by western blot in sWAT beige adipocytes treated as in b (n=3 per group). (d) Cellular respiration of sWAT beige adipocytes treated under the same conditions as in b and c (n=5 per group). (e) PKA activity in sWAT beige adipocytes treated without or with 50 mM Blebbistatin and without (0 min) or with 1, 2, 5, 10, 15, 20, and 25 minutes of 1 mM isoproterenol treatment (n=3 per group). (f) Lipolysis of sWAT beige adipocytes treated with 50 mM Blebbistatin and without or with 1, 2, 3, and 4 hours of 1 mM isoproterenol treatment as measured via free glycerol (n = 3 per group). (g) phospho and total HSL and CREB western blot of sWAT beige adipocytes treated as in e. (h) quantification of phospho to total CREB ratio of sWAT beige adipocytes as treated in g and e. Error bars represent mean ± SD. ∗p < 0.05, ∗∗p < 0.01. ∗∗∗p < 0.001, ∗∗∗∗ p < 0.0001

Canonically, b-AR agonism triggers thermogenic gene expression through the PKA-CREB pathway. However, to our surprise, blebbistatin had no effect on isoproterenol-stimulated PKA activity or lipolysis (Fig 4e, f) and western blots further confirmed that while blebbistatin treatment abrogated UCP1 expression, the same treatment did not alter isoproterenol-induced phosphorylation of hormone sensitive lipase (HSL) and cAMP response element-binding protein (CREB), two classical PKA effectors^48,49^ (Fig 4g, h). This was further confirmed with SVF-derived beige adipocytes where short-term blebbistatin did not impact isoproterenol-induced lipolysis (Supplementary Fig 6a). In white adipocytes, neither lipolysis induction (Supplementary Fig 6b) nor cellular respiration (Supplementary Fig 6c) was affected by blebbistatin treatment. These data indicate that while actomyosin activity is essential for β-adrenergic induction of UCP1 and thermogenic respiration, it does so without significant changes in canonical PKA activity and in a BeAT specific fashion, pointing to a novel non-canonical biomechanical signaling branch in this cell type.

### FAK mediates actomyosin-dependent events in b-adrenergic signaling

One of the best-described mechanisms through which actomyosin tension can activate biochemical signals is through force-based modulation of adhesive proteins at the cell-ECM interface^50–52^. One such mechanosensor is focal adhesion kinase (FAK), a cytoplasmic tyrosine kinase that becomes activated through autophosphorylation at tyrosine 397 (Y397)^50,51,53^. To determine whether FAK participates in the actomyosin-b-adrenergic axis in beige adipocytes, we first assessed its activation following b-adrenergic stimulation. Indeed, isoproterenol treatment rapidly induced FAK autophosphorylation at Y397 (Fig 5a). In contrast, isoproterenol treatment did not increase FAK phosphorylation in white adipocytes, indicating b-adrenergic driven FAK activation is beige-specific (Fig 5b, c). To test if this FAK activation is dependent on actomyosin contractility, we co-treated beige adipocytes with blebbistatin and isoproterenol. Blebbistatin abrogated isoproterenol-induced FAK phosphorylation (Fig 5d, e), demonstrating that FAK activation requires intact actomyosin tension. Similarly, in Myh9 KD cells, isoproterenol failed to induce FAK autophosphorylation (Supplementary Fig 7a, b), supporting that Myh9-mediated tension is required for b-adrenergic-induced FAK activation. Collectively, these results suggest that FAK functions as a tension sensor in beige adipocytes, transducing actomyosin contractility into downstream signaling.

**Figure 5:**
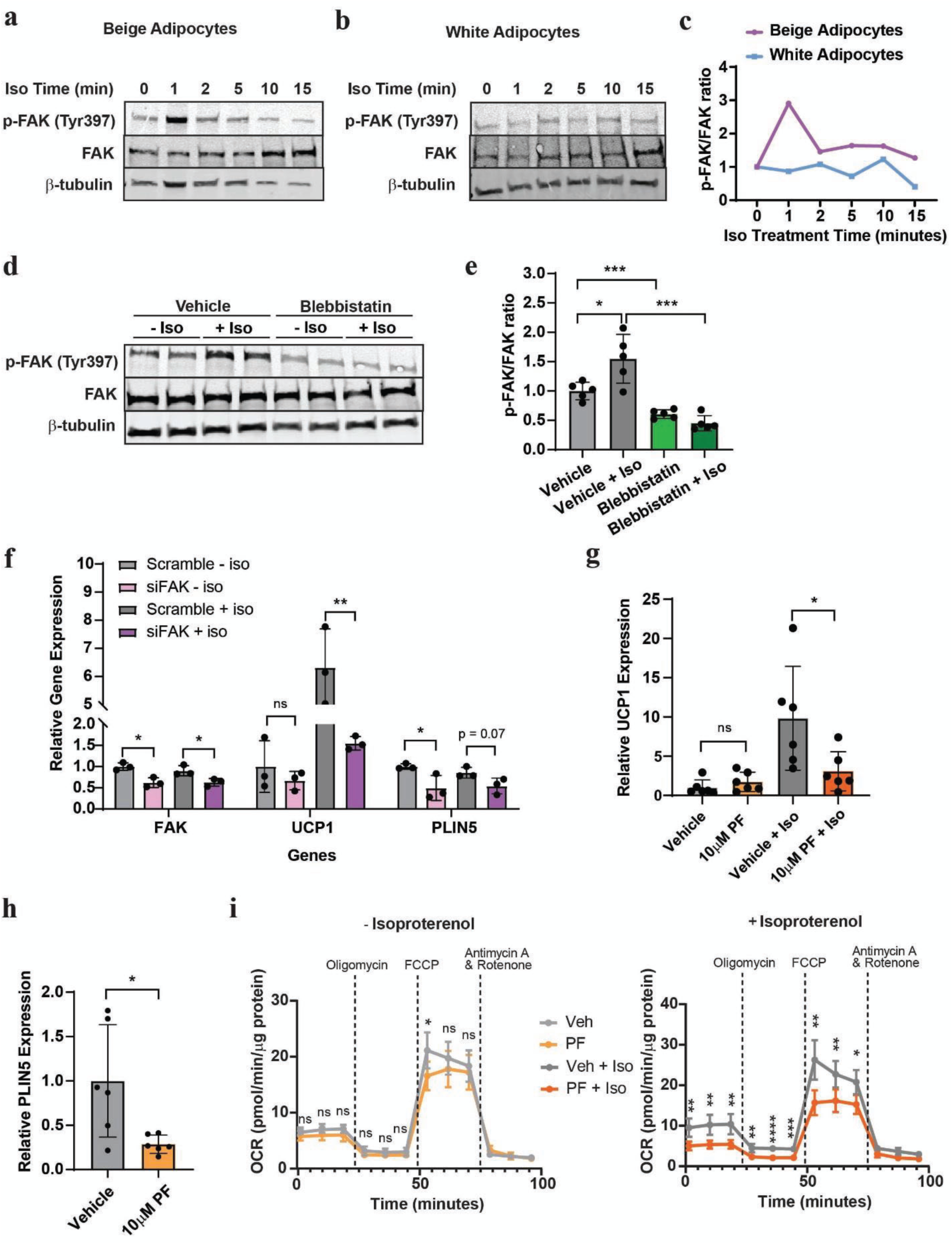
FAK transduces contractility signals to enhance UCP1 expression and thermogenesis. (a) Phospho-FAK (Y397) and total FAK level in sWAT beige adipocytes and (b) SVF-derived white adipocytes following 1, 2, 5, 10, and 15 minutes of 1 mM isoproterenol treatment, measured by western blot. (c) Quantification of a and b. (d) Phospho-FAK (Y397) and total FAK levels in sWAT beige adipocytes with or without 50 mM Blebbistatin and 1 mM isoproterenol (n=2 per group). (e) Quantification of phospho-FAK (Y397) and total FAK ratio in sWAT beige adipocytes as treated in a (n=5 per group). (f) Relative expression of FAK, UCP1, PLIN5 in sWAT beige adipocytes, treated with either scramble or FAK siRNA knockdown, and without or with 1mM isoproterenol measured by qPCR (n=3 per group). (g) UCP1 mRNA expression measured by qPCR in sWAT beige adipocytes treated overnight without (vehicle) or with 10 mM PF-573,228 and 1mM isoproterenol (n=6 per group). (h) PLIN5 mRNA expression measured by qPCR in sWAT beige adipocytes treated without or with 10 mM PF-573,228. (i) Cellular respiration of sWAT beige adipocytes treated without (vehicle) or with 10 mM PF-573,228 and 1mM isoproterenol for 2 hours (n=5 per group). Error bars represent mean ± SD. ∗: p < 0.05, ∗∗: p < 0.01. ∗∗∗: p < 0.001, ∗∗∗∗: p < 0.0001

We next examined whether FAK is required for b-adrenergic-induced UCP1 expression. siRNA-mediated knockdown of FAK in sWAT beige adipocytes significantly impaired isoproterenol-stimulated UCP1 expression, recapitulating the effects observed with Blebbistatin treatment and Myh9 KD when treated with isoproterenol (Fig 5f). Pharmacological inhibition of FAK using PF-573,228 (PF)^54,55^ effectively blocked FAK Y397 phosphorylation (Supplementary Fig 7c) and led to a significant reduction in isoproterenol-stimulated UCP1 expression (Fig 5g) and PLIN5 expression (Fig 5h), mirroring the phenotype of FAK siRNA KD. Same as UCP1 expression, isoproterenol-induced beige adipocyte basal and uncoupled respiration was also blunted when the cells are co-treated with PF (Fig 5i). Similar to activating actomyosin with EMD-57003, pharmacological activation of FAK with Zn27^56^ enhanced isoproterenol-stimulated UCP1 up-regulation (Supplementary Fig 7d).

## Discussion

Our study demonstrates that β-adrenergic stimulation in beige adipocytes, but not in white adipocytes, enhances actomyosin contraction-mediated intracellular tension and promotes rearrangement of actin filaments around lipid droplets, a process crucial for lipid droplet architecture. Loss of this biomechanical program through Myh9 depletion or acute myosin inhibition compromises bioenergetic gene expression, particularly UCP1 and PLIN5, and disrupts LD–mitochondria interactions. Notably, canonical PKA-mediated β-adrenergic signaling remains intact under actomyosin inhibition, implicating a parallel, tension-sensitive pathway mediated by FAK that is unique to beige adipocytes. Together, these findings identify a Myh9-driven mechanical signaling cascade that transcriptionally and functionally regulates thermogenesis in beige adipocytes.

BeAT and WAT are closely entwined in adipose tissues and are exposed to similar levels of b-AR signaling but respond differentially to this signal. In rodents, BeAT express all three β1, β2, and β3 adrenergic receptors^57^, with b3-AR being most enriched, whereas WAT primarily expresses β1 and β2-ARs with lower b3 abundance. These differences in receptor composition may modulate differential adrenergic responses, even though the downstream cAMP-PKA cascade is largely conserved between the two cell-types^58,59^. In our current study, we employed the non-selective β-adrenergic agonist isoproterenol, so it remains unclear whether all β-AR subtypes converge on the Myh9-mediated pathways or whether a particular receptor subtype predominantly initiates this pathway. Dissecting the contribution of each β-AR isoform using selective agonists and serial knockout/knockdown of each β-AR subtype would provide critical insights into this novel non-canonical pathway.

Besides differential expression of b-ARs, previous work on differential β-adrenergic responses in white versus beige adipocytes has focused largely on epigenetic mechanisms^60,61^. For example, Damal Villivalam et al. and Byun et al. independently reported that DNA demethylase TET1 acts as a BeAT-selective thermogenic repressor, which is suppressed under cold exposure to permit UCP1 induction^62,63^. Yet, a recent adipocyte epigenome atlas study revealed that while enhancer marks like H3K4me1 distinguish beige from brown adipocytes, murine beige and white adipocytes in iWAT are not inherently epigenetically distinct, likely due to common developmental progenitor^64^. These findings suggest that there exhibit mechanisms other than epigenetics that account for the two cell-types’ divergent responses to β-adrenergic stimulation. Our results provide an alternative explanation that biomechanical differences between the two cell-types might be the reason that the two cell-types exhibit differential response to the common b-adrenergic upstream.

FAK has been previously reported to be critical for beige adipocyte progenitor cells to respond to irisin via CD81 and integrin^65^. However, whether mature beige adipocytes require FAK activity was previously unknown. Here, our genetic and pharmacological loss-of-function studies on both myosin and FAK demonstrate that actomyosin contractility and FAK signaling are indispensable for b-adrenergic-induced thermogenic activation in mature beige adipocytes. Notably, gain-of-function experiments using pharmacological activators of actomyosin and FAK both only show enhanced UCP1 expression when co-stimulated with isoproterenol, suggesting that mechanical activation of actomyosin and FAK alone is insufficient to fully substitute for classical b-adrenergic signaling.

FAK activity is canonically coupled to integrin signaling at focal adhesion sites, linking intracellular actomyosin contractility with extracellular matrix (ECM) mechanical properties. Beyond typical hormonal regulation, emerging evidence highlights the importance of tissue microenvironmental components such as extracellular matrix (ECM) in shaping thermogenic adipocyte fate determination^66,67^. In obesity, WAT often undergoes fibrotic remodeling, leading to excessive ECM protein deposition and increased stiffness, which is negatively associated with beige adipocyte content.^68,69^. Adipocytes sense and adapt to ECM remodeling primarily through adhesion–mediated signaling ^52,70,71^ and a dynamic cytoskeletal network, in which the myosin family of actin-based motor proteins generates contractile forces against the tissue microenvironment via ECM adhesions^19,72^. Given that FAK activation involves both intracellular tension and ECM–integrin signaling, our findings provide a new aspect to the interplay between cell-intrinsic and ECM-derived mechanics. Interestingly, previous studies using in vitro hydrogel systems suggested that softening of ECM promoted white adipocyte differentiation^73^ where stiffening promoted beige and brown fat differentiation and metabolism^74–76^. Incorporating engineered biomaterial platforms with tunable biophysical properties may provide new opportunities to probe how ECM mechanics modulate Myh9–FAK signaling, with implications for designing therapeutic strategies that harness beige adipocytes to combat obesity and metabolic disease.

While identifying FAK as the mechano-sensor in beige adipocytes, other mechanotransducers such as MRTFA^77^ and YAP/TAZ^18,78,79^ were evaluated. MRTFA does not seem to drive the deficiency observed in our experiments, as blebbistatin-treated and Myh9-knockdown cells exhibited unchanged MRTFA nuclear translocation (Supplementary Fig 8). Meanwhile, although YAP/TAZ has been implicated in classical brown adipocyte function, our data and prior studies suggest it plays little role in beige adipocyte thermogenesis^78,79^. Instead, knockout of YAP/TAZ may promote adipocyte dedifferentiation through suppression of PPARG signaling, as reported by Choi et al^78^.

Which transcription factors mediate effects on thermogenesis downstream of FAK remains unclear. Some known FAK activated signals, such as STAT3 or ERK^80,81^, have been previously suggested to impact UCP1 expression in beige adipocytes^82–85^, but their precise roles in this context require further testing. On the other hand, while FAK inhibition disrupted β-adrenergic stimulation of UCP1 expression, it did not alter basal (unstimulated) UCP1 expression. In contrast, blebbistatin treatment or Myh9 KD showed reduced UCP1 levels both under basal and stimulated conditions. This distinction might suggest that FAK mediates acute β-adrenergic–induced UCP1 expression, whereas actomyosin tension more broadly contributes to beige adipocyte function, potentially via additional long-term signaling mechanisms that needs further investigation.

Beyond transcriptional control, our discovery that the Myh9–actin network stabilized LD–mitochondria contacts raises broader questions about how intracellular tension orchestrates organelle organization and bioenergetic reprogramming. Organelle trafficking plays an important role in physically connecting membrane-separated compartments, but the underlying mechanisms remain unclear. Our study provides direct evidence that, while PLIN5 has been implicated in anchoring LD–mitochondria contacts in previous studies, the Myh9-associated actomyosin network is the driving force behind this organelle organization in beige adipocytes. Interestingly, Myh9 KD and acute blebbistatin treatment produced divergent effects on lipolysis, possibly reflecting potential defects downstream of the chronic depletion of PLIN5 in the former versus short-term impairments of the latter. Future proteomic and lipidomic profiling of LD–mitochondria junctions, coupled with tension-sensor labeling, may reveal how biomechanical forces reshape metabolic connectivity in adipose tissue. The observation of dynamic “actin cage” formation also raises the question of whether this process is specific to unique subpopulations of LDs, potentially ensuring sufficient actin availability for the remaining LDs with varying lipolytic capacities. How actin is recruited to the LD surface and physically contributes to fatty acid trafficking from LDs into mitochondria remains unclear. Moving forward, isolating LDs, mitochondria, and actin filaments and studying their interactions *in vitro* may yield valuable insights into the biophysical basis of actin networks in organelle metabolism.

In summary, our work reveals that Myh9-dependent actomyosin tension is indispensable for β-adrenergic induction of UCP1 expression and for maintaining LD–mitochondria contacts in beige adipocytes. This tension-mediated pathway identifies a novel non-canonical branch of β-adrenergic signaling. These findings not only deepen our understanding of beige adipocyte biology but also suggest potential new targets for therapeutic activation of thermogenic fat to improve metabolic health.

## Methods

### Cell culture and differentiation

The immortalized murine beige adipocyte line (sWAT) is a generous gift from Shingo Kajimura^86^. Growth and differentiation of sWAT beige adipocytes and SVF derived beige and white adipocytes were described in our previous study^18^. Briefly, all cell lines were maintained in growth media consisting of DMEM supplemented with 10% fetal bovine serum (Equafetal) and 1% penicillin/streptomycin (Gibco). Differentiation was initiated once cells reached desired confluency.

To induce beige adipocyte differentiation, cells were cultured in induction media containing 5 mg/mL Insulin, 2 mg/mL Dexamethasone, 1 nM T3 (5 nM T3 for SVF derived beige adipocytes), 500 mM IBMX (Sigma), and 200 nM rosiglitazone (500 nM rosiglitazone for SVF derived beige adipocytes) upon reaching 95% confluency. After two days of induction, cells were switched to maintenance media with 5 mg/mL Insulin and 1 nM T3 (5 nM for SVF cells) for an additional four days. Cells were considered as fully differentiated at day 6 of differentiation.

For differentiation of white adipocytes, cells were induced once reached 100% confluent. Induction media for white cell differentiation contains 5 mg/mL Insulin, 2 mg/mL Dexamethasone, and 500 mM IBMX (Sigma). After two days of induction, cells were switched to maintenance media with 5 mg/mL Insulin after 2 days of differentiation induction. Cells were considered as mature at day 6 of differentiation.

All overnight treatments were conducted on day 6 of differentiation, and cells were harvested on day 7. Similarly, all short-term treatments are performed on differentiation day 7. Vendors and concentrations of compounds are listed below: Blebbistatin (50 mM, Cayman, #13013), EMD-57003 (10 mM, Sigma-Aldrich, #530657), PF-573,228 (10 mM, Sigma-Aldrich, #PZ0117), isoproterenol hydrochloride (1 mM, Sigma-Aldrich, #I6504).

### Animal and SVF extraction

All animal experiments were conducted under the guidelines and regulations by UC Berkeley Animal Care and Use Committee under protocol ID AUP-2019-05-12180-1. C57BL/6J mice were housed in an AAALAC-certified facility and monitored daily by the Office of Laboratory Animal Care (OLAC) veterinary lab staff for any health conditions during the experiments. Unless else noted, animals were fed a chow diet (LabDiet 5053) at 23 °C. For cold exposure experiments, 12 weeks old mice were cold exposed at 4°C for 24 hours using a Comprehensive Lab Animal Monitering System (CLAMS, Columbus Instruments).

To extract stromal vascular fraction (SVF), inguinal white adipose tissue (IWAT) was harvested from male and female mice following euthanasia by CO_2_ asphyxiation and cervical dislocation. The tissue was excised, rinsed in PBS, and minced using sterile scissors. Minced tissue was digested in 0.2% collagenase (Sigma-Aldrich, #C2-BIOC) dissolved in HBSS at 37°C for 50 minutes. Digestion was quenched by adding equal volume of DMEM/F12 supplemented with 10% FBS and 1% Pen/Strep. The suspension was then filtered through a 100 mm nylon mesh (Corning) and centrifuged at 500 rcf for 5 minutes. Cell pallet was resuspended in media and filtered through the 40 mm nylon mesh. After another centrifugation at 500 rcf for 5 minutes, cells were resuspended in erythrocyte lysis buffer () and incubated for 5 minutes at room temperature. Final centrifuge will be performed after incubation to remove the erythrocyte lysis buffer, and cells will be resuspended in DMEM/F12 supplemented with 10% FBS, 1% Pen/Strep, and 250 mg/mL amphotericin B and plated into 100 mm petri dishes.

### Plasmids, Lentivirus Production, Lentiviral Transduction

The pINDUCER lentiviral toolkit for inducible RNA interference 1 was used to generate Myh9 and Myh10 knockdown cell lines. pINDUCER11 miR-RUG (#44363) was purchased from Addgene. Myh9 and Myh10 miR30-Based shRNA vectors were derived as previously described. The shRNA top and bottom strands were annealed and then ligated into a pINDUCER11 vector that had been digested with EcoRI and XhoI. The ligated plasmid was transformed into One Shot TOP10 Chemically Competent E. coli (invitrogen C404010) and plasmid DNA was isolated using QIAprep Spin Miniprep Kit per manufacturer’s instructions. Lentiviruses were produced using Lipofectamine3000 (Invitrogen) according to the manufacturer’s instructions. Briefly, 293T cells were transfected with pINDUCER11, psPAX2 (a gift from Didier Trono, Addgene plasmid # 12260; RRID:Addgene_12260), and pMD2.G (gift from Didier Trono, Addgene plasmid # 12259 ; RRID:Addgene_12259) in Lipofectamine3000. Supernatants were harvested at 24h and 48h later, clarified, filtered through a 0.45-μm filter and stored at -80°C. sWAT and sBAT preadipocytes were transduced with lentivirus and 10ug/ml of polybrene. After 2 days post transduction, cells positive for GFP were selected to generate inducible shRNA stable cell lines.

Myh9 shRNA oligos:

Top Strand 5′-

TCGAGAAGGTATATTGCTGTTGACAGTGAGCGCTACCCTTTGAGAATCTGATACTAGTGAAGCCACAGATGTAGTATCAGATTCTCAAAGGGTAGTGCCTACTGCCTCGG-3′)

Bottom Strand

5’AATTCCGAGGCAGTAGGCACTACCCTTTGAGAATCTGATACTACATCTGTGGCTTCACTAGTATCAGATTCTCAAAGGGTAGCGCTCACTGTCAACAGCAATATACCTTC-3’)

Myh10 shRNA oligos:

Top Strand 5′-

TCGAGAAGGTATATTGCTGTTGACAGTGAGCGCCCTCCACAAGACATGCGTATTTAGT

GAAGCCACAGATGTAAATACGCATGTCTTGTGGAGGGTGCCTACTGCCTCGG-3’)

Bottom Strand

5’AATTCCGAGGCAGTAGGCACCCTCCACAAGACATGCGTATTTACATCTGTGGCTTCACTAAATACGCATGTCTTGTGGAGGGCGCTCACTGTCAACAGCAATATACCTTC-3’)

PLIN5 overexpression (OE) cell lines were generated using Genscript GenEZ™ ORF clones with pGenLenti backbone (Plin5_OMu18852C_pGenLenti). pLenti Lifeact-iRFP670 BlastR was a gift from Ghassan Mouneimne (Addgene plasmid # 84385 ; http://n2t.net/addgene:84385 ; RRID:Addgene_84385^87^) and was used to generated cells with fluorescent F-actin labels. Subsequent lentiviral production and transduction were performed as described earlier in the section. Puromycin and blasticidin selection were initiated 24 h post infection accordingly for PLIN5 OE and Liface-iRFP670 cell lines.

### siRNA-mediated knockdown of target genes

Scramble or predesigned siRNA targeting mouse Myh9, Myh10, FAK and PLIN5(MISSION siRNA, Millipore Sigma) was mixed with lipofectamine RNAi MAX reagent (thermo fisher) and applied on differentiation day 1 and day 4 of cultured adipocytes overnight (20 pmol of siRNA per one well of 12-well plate). siRNA universal negative controls (MISSION siRNA, Millipore Sigma) were used as scramble controls for siRNA knockdown experiments.

### RNA isolation and qRT-PCR

mRNA was isolated from tissues or in vitro cultures with TRIzol reagent (invitrogen), and purified using Monarch total RNA miniprep kit (NEB). cDNA was synthesized using Maxima First Strand cDNA synthesis kit (K1672, thermo scientific), and 10 ng cDNA was used for qPCR on a QuantStudio 5 real-time PCR system with TaqMan Universal Master Mix II and validated PrimeTime primer probe sets (Integrated DNA Technologies). The DDCT method was used to comparatively assess mRNA quantity. Probes for qRT-PCR are listed in Supplementary Table 1.

### Protein extraction and western blotting

Protein was extracted from tissues or in vitro cultures with RIPA buffer (Invitrogen) supplemented with 1% Halt proteinase and phosphatase inhibitor cocktail (Protein Biology). Protein quantity was measured using Pierce BCA reagent kit (Thermo Scientific). Protein lysates were either used immediately or stored at -80°C for future use. Equal protein concentrations were mixed with a 6x Laemmli buffer containing 2-Mercaptoethanol and boiled at 95°C for 4 minutes. Samples were resolved on Bio-RAD Protean-TGX gels and transferred onto nitrocellulose membrane using Bio-Rad Transblot turbo. The membranes were blocked with 5% non-fat milk dissolved in 0.1% TBST for one hour and then incubated with primary antibodies for overnight. Secondary antibodies were incubated for 1 hour, and blots were imaged using Odyssey Imaging System. Primary and secondary antibodies used were listed in Supplementary Table 1.

### PKA assay

To investigate b-AR canonical downstream, PKA activity was measured with ELISA based PKA Calorimetric activity measurement kit from Invitrogen (Invitrogen, EIAPKA). Cells are treated without (0 min) or with 1mM isoproterenol for 1, 2, 5, 10, 15, 20, 25 minutes, and cell lysates are harvested and tested following the kit protocol. Measured PKA activities are normalized to the non-treated control group.

### Free glycerol assay

Glycerol release assay was performed using Caymen Glycerol Cell-Based Assay Kit (10011725), and free glycerol reagent from Sigma-Aldrich (F6428). Briefly, cells were differentiated into beige adipocytes in 96 well plates. On day 7, cell media was replaced with 50μl adipolysis buffer (0.5% BSA in DMEM) for 4 hours. 25μl media from each well was then collected and mixed with 100 μl of reconstituted Free Glycerol Assay Reagent per well. After incubation for 15 minutes at room temperature, the absorbance was read at 540 nm on a SpectraMax i3 plate reader and the data was normalized to standard curves and plotted.

### In vitro respirometry (seahorse assay)

Oxygen consumption rate (OCR) was measured in beige adipocytes cultured and differentiated in a 24-well Seahorse plate using the Seahorse XFe24. Prior to experiments, cells were incubated in the XF assay medium supplemented with 25 mM glucose, 5 mM sodium pyruvate, and 2 mM glutamine in a 37 °C CO2 free incubator for an hour. Cells were subjected to the mitochondrial stress test by sequentially adding oligomycin (1 μM), FCCP (1 μM), and antimycin/rotenone mix (1 μM/1 μM). Measurements were normalized on protein content performed by BCA assay.

### Quencher-based real time BODIPY-fatty acid uptake assay

The real-time fatty acid uptake assay was based on a previously published assay^88^. Cells were plated in 96-well clear bottom tissue culture plates and differentiated toward mature adipocytes. Cell media was replaced with uptake solution containing 1mM trypan blue, 3.5 g/L glucose, 2μM BODIPY-fatty acid, and 0.1% bovine serum albumin in Hank’s buffered saline solution. The cells were immediately placed into a SpectraMax i3 plate reader and the fluorescence was read at an excitation of 488 nm and emission of 515 nm every minute for 90 minutes. The steady-state fluorescence at the end was plotted to show the overall differences.

### Organelle staining for live cells and immunofluorescence staining for fixed samples

For live cell imaging, the cells were plated onto 96 well polymer dishes with tissue culture treatment (Cellvis P96-1.5P) and stained with organelle dyes per manufacturer’s instructions. Mitochondria: MitoBright LT Green (Dojindo, MT10), MitoBright ROS Deep Red (Dojindo, MT16-10), PKmito ORANGE (Cytoskeleton, CY-SC053); Lipid droplets: Lipidblue (Dojindo, LD01-10), BODIPY™ 493/503 (Thermo Fischer D3922), F-actin: SPY650-actin (Spirochrome SC505). Dishes were rinsed gently with PBS three times and replaced with phenol red free DMEM 25mM HEPES buffer imaging media. Images were acquired on Zeiss LSM 980 with Airyscan at the UC Berkeley CRL Molecular Imaging Center.

For fixed samples, cells were fixed with 4% PFA for 10 minutes at room temperature followed by 3x washes with PBS. The cells were then permeabilized with 0.05% Triton X in PBS for 15 minutes at room temperature. After permeabilization, the cells were incubated in blocking buffer (1% BSA in PBS) for 1 hour at room temperature. Cells were then incubated with primary antibodies diluted in 0.05% Triton X in 1% BSA in PBS for overnight at 4°C. Following primary incubation the cells were washed 3x with blocking buffer and incubated with secondary dyes diluted in blocking buffer for 1 hour at room temperature. Following secondary incubation, the cells were washed 3x with and ready for imaging sessions. Primary antibodies (dilution, company, catalog number): anti-Myh9 (1:100, Proteintech, 11128-1-AP), anti-MyhIIB (1:100, Abcame, ab300647), anti-TOM20 (1:300, Sigma-Aldrich, WH0009804M1). Samples were incubated with goat anti rabbit AlexaFluor 647 secondary antibodies, donkey anti rat AlexaFluor antibody 555 as well as Alexa Fluor 488, 594, 647 Phalloidin, and 4’,6-diamidino-2-phenylindole (DAPI) from Thermo Fischer Scientific.

### Microscopy and image analysis

Images were acquired on Nikon AX/AX R with NSPARC Confocal, Zeiss/Yokagawa Spinning Disk, Zeiss LSM880 FCS, Zeiss Elyra 7 Lattice SIM-squared, and Zeiss LSM 980 NLO with Airyscan system. EC Plan-Neofluar 40x/1.30 Oil DIC M27 and Plan-Apochromat 63x/1.40 Oil DIC M27 were used as objectives for various imaging experiment set ups. Fluorescence channels were set up using smart set up options in Zeiss Zen software and customized for single acquisition, time course images, or stitched tile scanning. For Airyscan images, images were further processed within Zen software for “Airyscan processing” allowing for pixel reassignment and deconvolution to generate the final super-resolution images. For lattice-SIM microscopy, images were further processed by Lattice-SIM^2^ reconstruction processing tool of the ZEN software to increase the resolution of the microscopy data. At least 9 images were obtained for each sample and at least three samples from each treatment group were used for analysis. Overnight time lapse images were acquired on Perkin Elmer Opera Phenix with 40x air objective lens. 10 randomly selected positions per 96 well plate was imaged at 15-minute intervals for 12 hours.

All images were analyzed in ImageJ/Fiji. For the quantification of LD sizes, Otsu threshold method was used to identify a region of interest (ROI) mask for individual LD. “Watershed” function was applied to further separate LDs that were in contact with each other. ROI masks were then analyzed to obtain size information with a cut off size of at least 1μm^2^. For intensity comparison of mitochondria ROS, mean background signals were calculated and subtracted from raw images. Positive pixels were then threshold to obtain mitochondria masks. Integrated intensity of each mitochondria masks was measured and normalized to the average intensity of vehicle control cell samples. For LD-mitochondria contact measurements, both LD and mitochondria ROI masks per image were isolated. Then mitochondria masks were overlaid with LD masks to generate LD-mitochondria contact masks. The area of contact masks and LD masks were calculated and divided to obtain % LD-mitochondria contact areas.

### Laser nanosurgery

All experiments were performed on a Zeiss LSM 980 confocal microscope equipped with a mode-locked MaiTai Ti:sapphire femtosecond laser (Spectra Physics). Protocols were adapted based on previously published laser nanosurgery protocols for single stress fiber ablation^89^. Cells were stained with live organelle dye for LD (lipidblue) and F-actin (SPY650-actin) as described previously. Confocal fluorescence images were obtained by illuminating samples with 405 and 639 laser lines. Simultaneously, a narrow beam (area 1 μm^2^) from the femtosecond laser was tuned to 760 nm for 10 iterations to photo irradiate actin signals near LDs to sever the structures. Sequential confocal images were obtained for up to a total of 30 seconds at 1s intervals post laser ablation to track LD size dynamics. Using ImageJ/Fiji, LD masks were isolated at each point and the coordinates of two merging LDs were measured via “center of mass” function. The distance of LDs traveled per second post ablation was quantified via 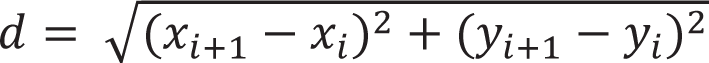

### Cytosolic Ca^2+^ imaging

Cells were incubated with 3 μM Fluo-4 AM (Invitrogen, F14201) and 0.1% Pluronic F-127 (Invitrogen, P3000MP) in Hanks’ balanced salt solution (HBSS) for approximately 60 minutes at room temperature in the dark. After incubation, cells were washed twice with HBSS and maintained in HBSS during imaging. Live-cell calcium imaging was performed using a Zeiss/Yokogawa spinning-disk confocal microscope with a 63× oil-immersion objective. Time-lapse images were acquired every 5 seconds for a total duration of 25 minutes. Isoproterenol (1 μM final concentration) was added at the 1-minute time point. Image sequences were analyzed using ImageJ. Regions of interest (ROIs) corresponding to individual cells were manually defined and applied across all time frames. Mean fluorescence intensity within each ROI was measured at each time point to quantify cytosolic Fluo-4 AM fluorescence.

### Cellular contraction imaging

For visualization of actin dynamics during contraction, cells were incubated with SPY650-FastAct (Cytoskeleton, CY-SC505) diluted in HBSS at 37 °C for approximately 2 hours before imaging. Time-lapse imaging was performed using a Zeiss/Yokogawa spinning-disk confocal microscope with a 63x oil objective similar to cytosolic calcium imaging. One minute after acquisition began, cells were treated with 1 μM isoproterenol (final concentration). To quantify cellular contraction, individual cells were manually outlined as ROIs, and cell areas were measured at the start and at the end of the recording. Relative cell area was calculated as the ratio of the post-treatment area to the pre-treatment area.

### AFM

Freshly isolated mouse BAT and iWAT pieces were rinsed with DMEM (Agilent) supplemented with 25 mM HEPES. The tissues were thoroughly cleaned of non-adipose material and cut into ∼1mm cube pieces. For better adhesion, 0.5% agarose diluted in DMEM/F12 was heated to coat the center of a 50×9mm style polystyrene petri dish (Falcon, 351006). Fresh tissue samples were placed in the center of the dish and allowed solidification of agarose to happen to immobilize the tissue samples. Cell samples were seeded on glass coverslips coated with 0.5μg/ml collagen and differentiated as mature beige adipocytes. Each coverslip was then transferred to and glued to the center of petri dishes with vacuum grease and immersed with DMEM media supplemented with 25mM HEPES buffer.

Tissue and cell stiffness measurements were performed by AFM (Asylum Research, Oxford Instruments) as previously described^90^. Briefly, AFM silicon nitride probes with triangular silicon nitride cantilevers and gold coating (PNP-TR, spring constant (k) ≈ 0.08 N/m; Nanoworld) were calibrated by the thermal method. Contact mode force measurements were carried out with a trigger force of 3 nN and a scan rate of 0.25 μm/s. Force maps of 8×8 force curves were taken in multiple randomly selected 10×10 μm scan grid for each sample. Elastic modulus was calculated from mean force curves using the system software by fitting the Hertz model.

### Transmission electron microscopy

Cells were grown on 35mm MatTek glass-bottom dishes (#P35G-1.5-14-C MatTek Corp., Ashland, MA, USA) and fixed with 1 mL of fixative media for a minimum of 20 min. (Fixative: 2.5% glutaraldehyde and 2.5% paraformaldehyde in 0.1M sodium cacodylate buffer, pH 7.4 (EMS, Hatfield, PA, USA). Samples were rinsed (3×; 5 min, Room Temperature) in 0.1M sodium cacodylate buffer, pH 7.2, and immersed in 1% osmium tetroxide with 1.6% potassium ferricyanide in 0.1M sodium cacodylate buffer for 30 minutes. Samples were rinsed (3×; 5 min, RT) in buffer and briefly washed with distilled water (1×; 1 min, RT), then subjected to an ascending ethanol gradient followed by pure ethanol. Samples were progressively infiltrated (using ethanol as the solvent) with Epon resin (EMS, Hatfield, PA, USA) and polymerized at 60 C for 24-48 hours. Care was taken to ensure only a thin amount of resin remained within the glass bottom dishes to enable the best possible chance for separation of the glass coverslip. Following polymerization, the glass coverslips were removed using ultra-thin Personna razor blades (EMS, Hatfield, PA, USA) and liquid nitrogen exposure, as needed. Regions of interest were cut and mounted on a blank resin block with cyanoacrylate glue for sectioning. Thin sections (80 nm) were cut using a Leica UC6 ultramicrotome (Leica, Wetzlar, Germany) from the surface of the block and collected onto formvar-coated 50 mesh grids. The grids were post-stained with 2% uranyl acetate followed by Reynold’s lead citrate, for 5 min each. The sections were imaged using a FEI Tecnai 12 120kV TEM (FEI, Hillsboro, OR, USA) and data recorded using a Gatan Rio 16 CMOS with Gatan Microscopy Suite software (Gatan Inc., Pleasanton, CA, USA). Organelle was hand traced using ImageJ software using the freehand tool.

### Bulk RNA-seq

Murine beige adipocytes (sWAT) were differentiated for 7 days with or without Myh9 knockdown and n=3 independent biological replicates were pooled to generate total RNA samples. Sequencing libraries were constructed from mRNA using KAPA mRNA HyperPrep Kit (KK8580) and NEXFlex barcoded adaptors (Bioo Scientific). High-throughput sequencing was performed using a HiSeq 2500 instrument (Illumina) at the UC Berkely QB3 Core. Raw reads were mapped using STAR60 against mouse (mm10) genome. Subsequent analyses of differential gene expression were carried out using Python (3.10.13) with the methods described below.

### Differential Expression Analysis

RNA-seq gene counts were first filtered using a minimum total counts of 100 to filter out low-count genes prior to standard differential expression pipeline. Differential expression analysis (DEA) was conducted with pyDESeq2^91^, a python implementation of DESeq2. Differentially expressed genes (DEGs) were identified with Wald test using pyDESeq2 (with threshold false discovery rate (FDR/adjusted p-value) ≤ 0.05; absolute log2 fold change ≥ 1) after multiple testing correction by Benjamini–Hochberg method. Visualization of DEGs is performed with Matplotlib (3.8.3), Seaborn (0.13.0), and Scipy (1.11.3).

Top 50 up and down differentially expressed genes were referenced with The National Center for Biotechnology Information (NCBI) database for the subcellular distribution identification.

### Enrichment Analysis

To gain thorough understanding of biological functions that have be altered, we performed enrichment analysis using SRplot^92^ based on the DEGs extracted from differential expression analysis and sorted by log2 fold change. After Benjamini–Hochberg correction, terms and pathways that were significantly enriched were identified (with threshold adjusted p-value ≤ 0.05). Significantly enriched terms were selected based on biological functions and plotted in Python using Matplotlib (3.8.3) and Seaborn (0.13.0).

### Statistics

All experiments have been reproduced at least 3 times if not other stated in figure legends, with biological replicates derived from independent samples (n). In all graphs, dots represent the values of each biological replicate and mean values were calculated from independent samples. All results from the tissue culture condition were obtained from at least 3 biological replicates. All results were statistically analyzed and plotted using Prism (GraphPad). Two-tailed student t-test with assumption of equal variance was used to compare the difference between two groups and one-way analysis of variance (ANOVA) followed by Tukey’s multiple comparisons test was used for more groups. Non-paired comparison is used in most cases excepting noted pairwise. P-value <0.05 was evaluated as significant (*), <0.005 as highly significant (**). Results were presented in mean values. Error bars represent standard deviation of the mean (SD) in each group unless noted in the figure captions.

Graphics:

All graphic illustrations were created using BioRender.com. All Figures were arranged using Adobe Illustrator.

## Data Availability

List of imaging reagents, qPCR probes, and public datasets are provided in Supplementary Tables 1. Transcriptomic raw data is available on public repository, Gene Expression Omnibus, under GSE311191.

## Acknowledgement

This study was supported by NIH grant R01DK118940-03 and R01DK141135-01. Y. He and L. Ling received fellowship from the California Institute of Regenerative Medicine (CIRM) EDUC4-12790. Airyscan confocal and two-photon imaging experiments were conducted at the CRL Molecular Imaging Center, RRID:SCR_017852, supported by NIH S10OD025063.

We would like to thank Holly Aaron and Luis Alvarez for their microscopy advice and support. Confocal and super-resolution microscopy was performed at RCNR Biological Imaging Facility and was supported in part from 1S10RR026866-01 and NIH 1S10OD018136-01, and we thank Dr. Denise Schichnes, and Dr. Steve Ruzin for their support. Research reported in this publication using Perkin Elmer Opera Phenix was supported by the Office of the Director, National Institutes of Health, under Award Number S10OD021828. We thank Mary West at the High-Throughput Screening Facility (HTSF) at UC Berkeley for her training of this machine. We also want to thank Dr. Elizabeth Obrien and Dr. Thomas Kenney at Nikon Instruments for their support in using Nikon AX/AX R system. TEM work was supported via NIH S10 Grant #S10OD030258-01 and we would like to thank Dr. Danielle Jorgens and Reena Zalpuri at the UC Berkeley Electron Microscope Laboratory for advice and assistance in electron microscopy sample preparation and data collection.

## Author Contributions

Y.H., L.L., and G.D. performed all experiments and data analysis. N. K., Z. V., E. C., I. L., I. C., V. T. facilitated Y.H. with experiments and data analysis. Y.H., L.L, S.K., and A.S designed, supervised the experiments and wrote the paper. All authors were involved with editing the paper and discussion of results. G.D. initially identified the phenotypes, performed siRNA screening of Myh genes, and generated Myh9 and Myh10 knockdown cell lines. Y.H. and G.D. carried out gene expression and respiration analyses in Myh9 KD and pharmacologically treated cells. Y.H. confirmed actomyosin contraction and upstream calcium influx, analyzed public datasets to compare beige and white adipocytes, validated canonical adrenergic signaling, and identified FAK that links actomyosin activity to UCP1 expression. L.L. conducted AFM measurements, established the relationship between F-actin, Myh9, and lipid droplet dynamics, and identified lipid metabolism alterations. L.L and G.D. performed EM studies on Myh9 KD cells, with L.L. analyzing altered LD–mitochondrial interactions and confirming PLIN5’s role via PLIN5 overexpression experiments.

