## Supplemental figures for "Bioenergetic responses to β-adrenergic stimulation in beige adipocyte depend on actomyosin driven forces"

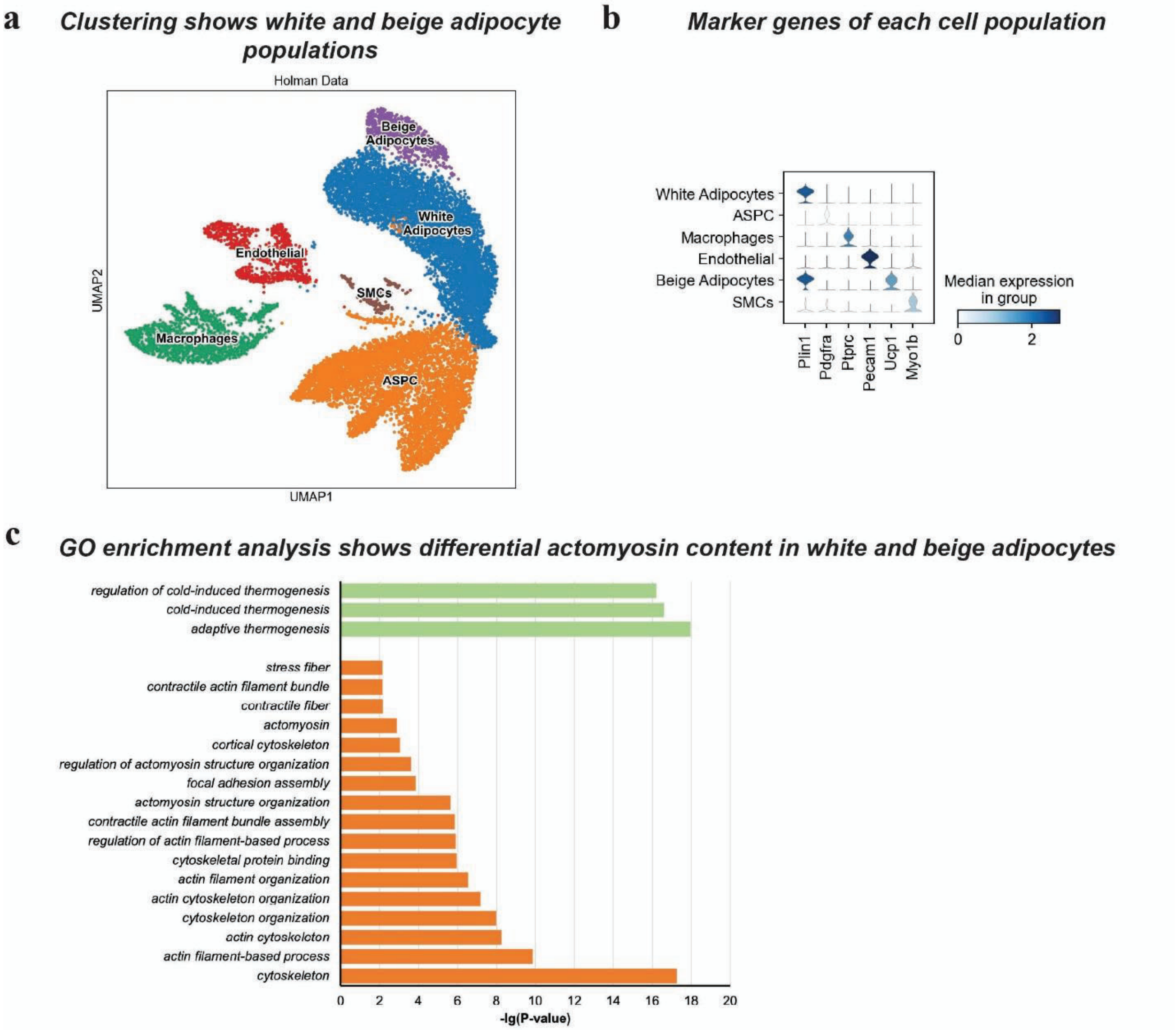


**Supplementary Figure 1. Public snRNA-seq reveals differential actomyosin gene expression profile in white and beige adipocytes** (a) Clustering reveals distinct beige and white adipocytes. (b) Marker genes of each major cell population identified in a. (c) Enrichment analysis of GO Biological Process of up-regulated differentially expressed genes in beige adipocyte population when compared to white adipocytes reveal enriched stress fiber, actomyosin, and focal adhesion related terms (dark orange) along with thermogenesis related terms (green). Differentially expressed genes of cortical actin, ECM, and stress fiber related genes in IWAT of cold exposed mice and room temperature housed control.

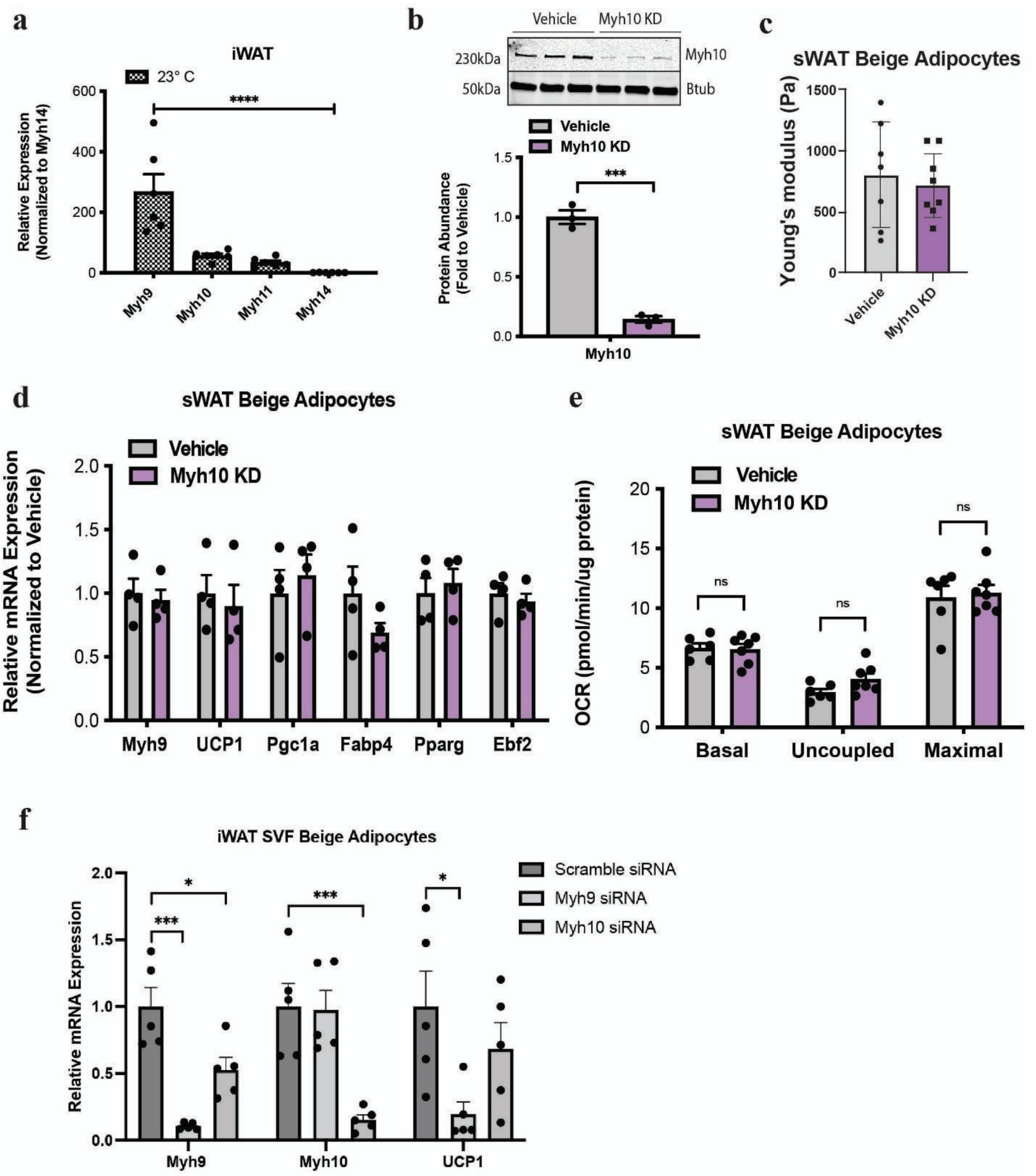


**Supplementary Figure 2.** (a) Relative expression of myosin heavy chains (Myh) in mouse inguinal white adipose tissue (IWAT) as measured via qPCR (n=6 mice per group). (b) Western blot measurement of Myh10 level in Myh10 KD cells and un-induced vehicle control (n=3 per group) (c) AFM measurements of cellular tension for Myh10 shKD cells vs vehicle control. (d) UCP1, Pgc1a, Fabp4, Pparg, Ebf2 mRNA expression measured by qPCR in inducible Myh10 KD sWAT beige adipocytes without (vehicle) or with induction of Myh10 shRNA expression for 7 days during differentiation (n=3 per group). (e) Cellular respiration of inducible Myh10 KD sWAT beige adipocytes without (vehicle) or with induction of Myh10 shRNA expression for 7 days during differentiation (n=5 per group). (f) Relative expression by qPCR of Myh9, Myh10, and UCP1 in mature beige adipocytes differentiated from IWAT derived SVF cells, treated with either scramble or with Myh9/Myh10 siRNA (n=5 per group).

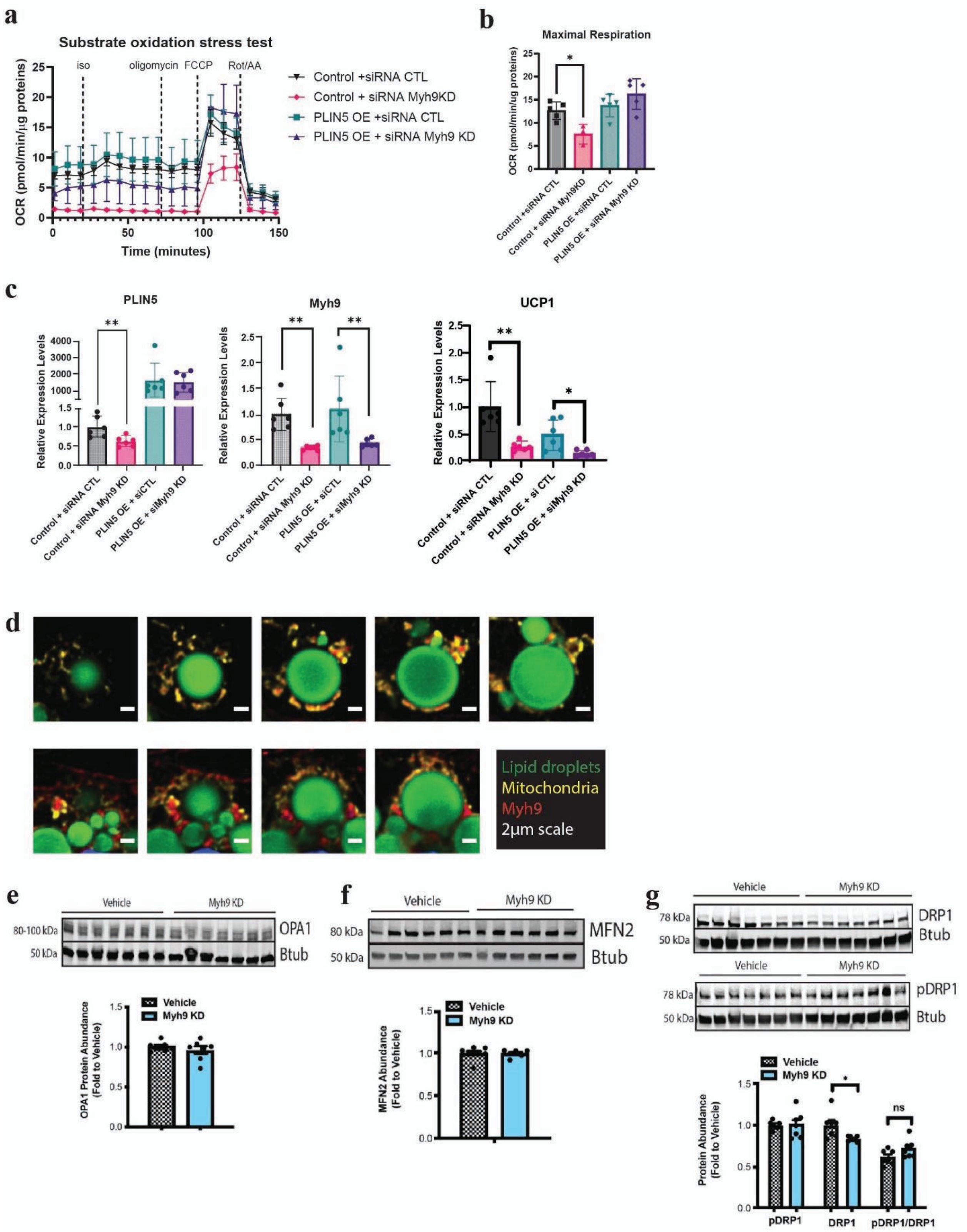


**Supplementary Figure 3.** (a) Cellular respiration of PLIN5 OE and regular beige adipocytes treated with siRNA control (CTL) or Myh9 KD (n=3-5 per group). (b) Quantification of maximal respiration as measured in (a). (c) Gene expression in PLIN5 OE and regular beige adipocytes treated with siRNA control (CTL) or Myh9 KD. (d) Immunofluorescence images showing subcellular distribution of Myh9 (red), mitochondria (yellow), and LD (green). 2µm scale (e-g) Western blot of three mitochondria fission/fusion proteins (OPA1, MFN2, and DRP1) from vehicle vs Myh9KD beige adipocytes, n=6-7 independent biological replicates.

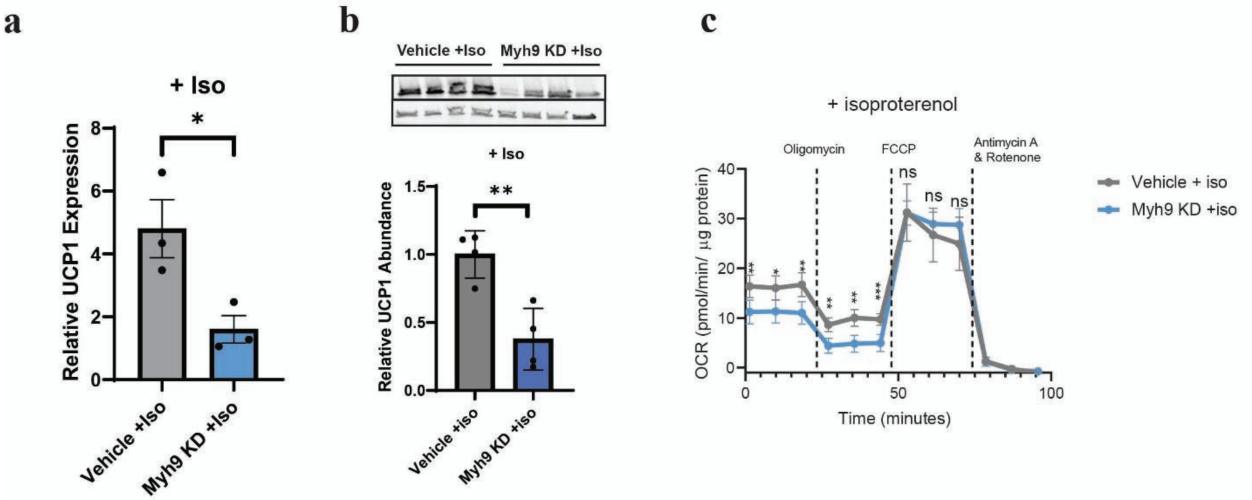


**Supplementary Figure 4. Acute ß-AR signaling blocked by Myh9 KD.** (a) UCP1 mRNA and (b) protein expression measured by qPCR and western blot in inducible Myh9 KD sWAT beige adipocytes without (vehicle) or with induction of Myh9 shRNA expression for 7 days during differentiation, followed by overnight treatment with 1mM isoproterenol. (c) Cellular respiration of inducible Myh9 KD sWAT beige adipocytes without (vehicle) or with induction of Myh9 shRNA expression for 7 days during differentiation, followed by a 2-hour treatment with 1mM isoproterenol (n=5 per group).

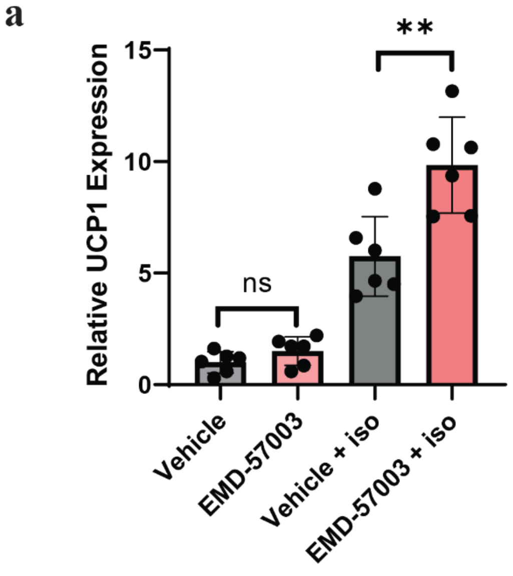


**Supplementary Figure 5. EMD-57003 amplifies b-adrenergic stimulation of UCP1 expression and uncoupled respiration.** (a) UCP1 mRNA expression measured by qPCR in sWAT beige adipocytes pre-treated overnight without (vehicle) or with 10 mM EMD-57003, followed by a 2-hour treatment with or without 1mM isoproterenol (n=6 per group).

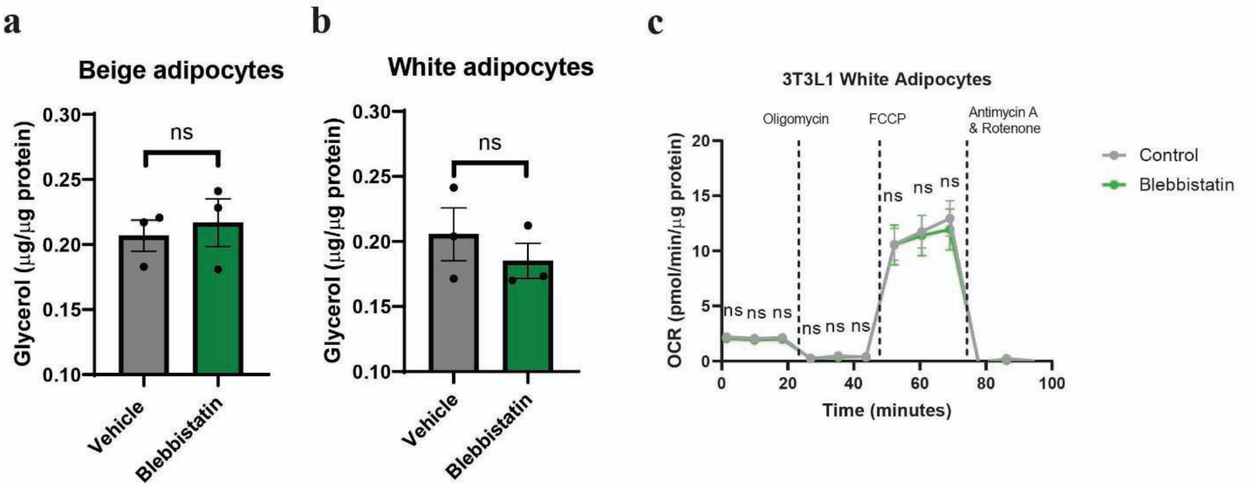


**Supplementary Figure 6. White adipocyte supplementary data.** (a) Lipolysis of SVF derived beige and (b) white adipocytes after 4 hours of 1 mM isoproterenol treatment with or without 50 mM Blebbistatin (n = 3 per group). (c) Cellular respiration of 3T3L1 white adipocyte treated overnight without or with 50 mM Blebbistatin.

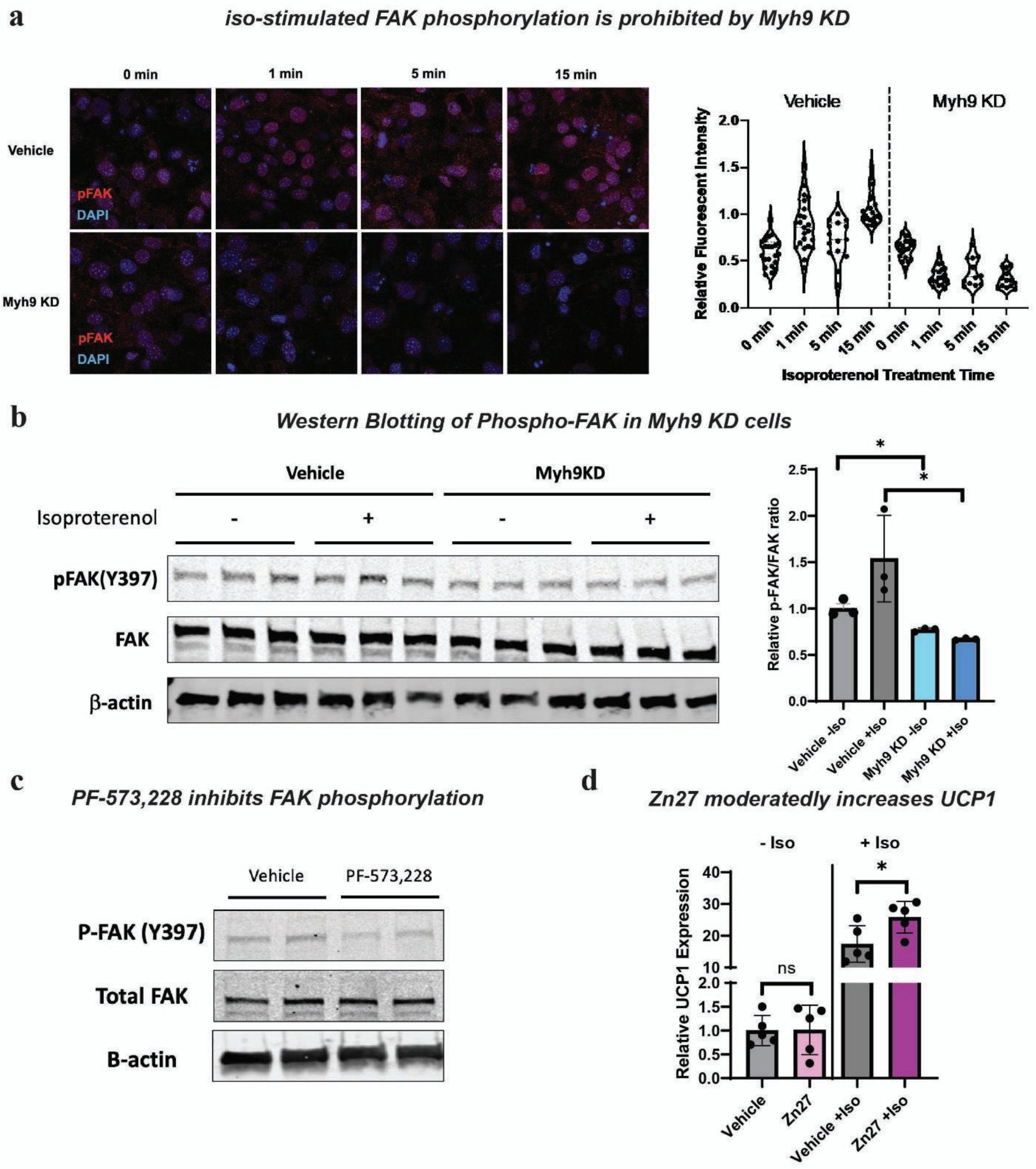


**Supplementary Figure 7. FAK supplementary data.** (a) Immunofluorescent imaging of phospho-FAK (Y397) (red) and DAPI (blue) in inducible Myh9 KD sWAT beige adipocytes, treated without or with induction of Myh9 shRNA expression and 1 mM isoproterenol. (b) Phospho-FAK (Y397) and total FAK level in Myh9 KD sWAT beige adipocytes following, treated without or with Dox induction during differentiation and 1 mM isoproterenol. (c) Phospho-FAK (Y397) and total FAK level in sWAT beige adipocytes following 10 mM PF-573,228 treatment. (d) UCP1 mRNA expression measured by qPCR in sWAT beige adipocytes treated overnight without (vehicle) or with 10 mM Zn27 and 1mM isoproterenol (n=5 per group)

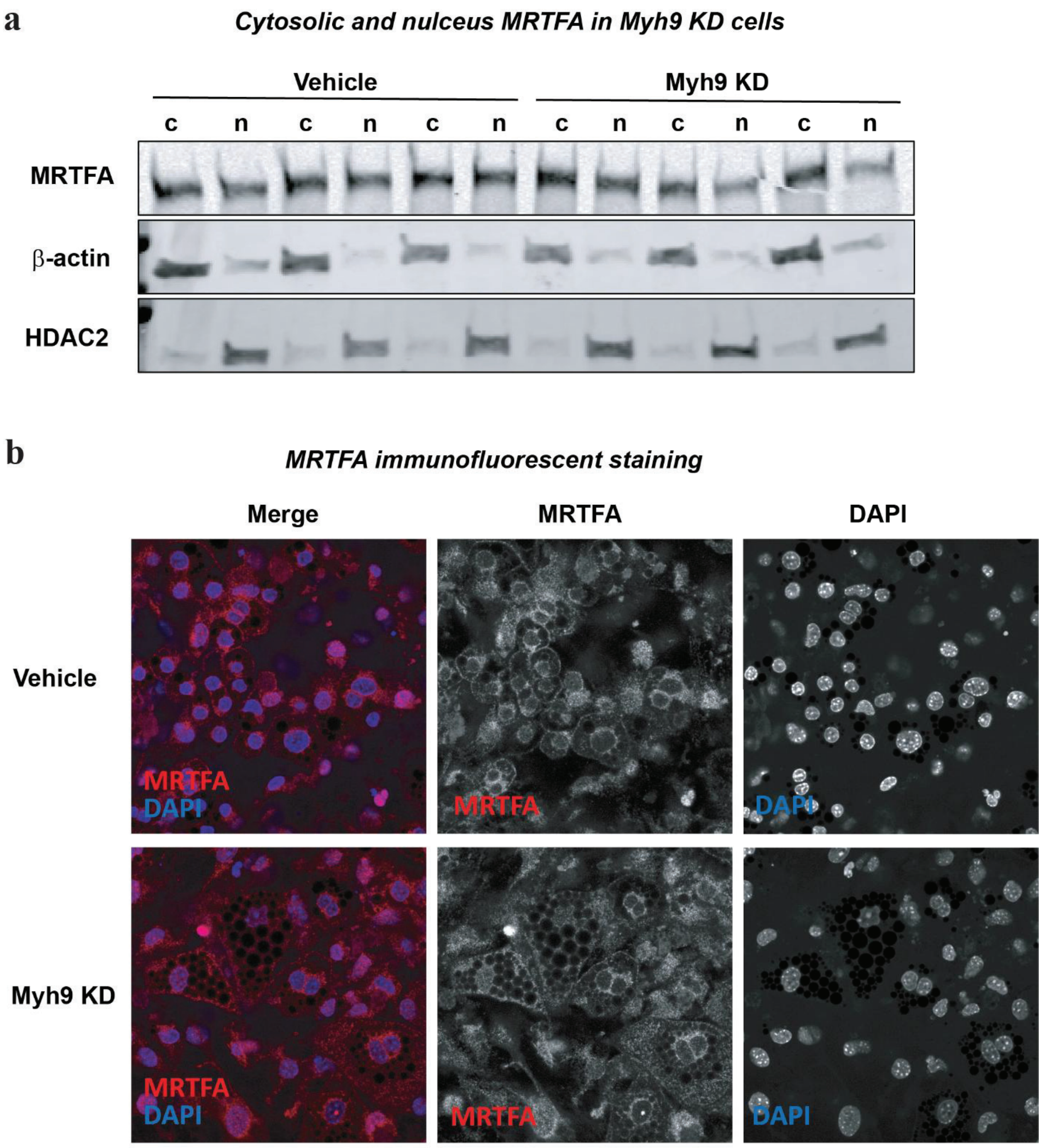


**Supplemental figure 8.** (a) Western blot of cytosolic and nucleus MRTFA. **b.** Immunofluorescent staining of MRTFA in Vehicle and Myh9 KD cells.

| Reagents | Source | Identifier/catalog number | Concentrations/Dilution factors |
|---|---|---|---|
| rt-qPCR probes | | | |
| Ucp1 | Integrated DNA Technologies | Mm.PT.58.7088262 | n/a |
| Myh9 | Integrated DNA Technologies | Mm.PT.58.13928218 | n/a |
| Myh10 | Integrated DNA Technologies | Mm.PT.58.30923223 | n/a |
| Myh11 | Integrated DNA Technologies | Mm.PT.58.9236105 | n/a |
| Myh14 | Integrated DNA Technologies | Mm.PT.58.41356163 | n/a |
| Fabp4 | Integrated DNA Technologies | Mm.PT.58.43866459 | n/a |
| Glut4 | Integrated DNA Technologies | Mm.PT.58.9683859 | n/a |
| Pnpla2 | Integrated DNA Technologies | Mm.PT.56a.13182140 | n/a |
| Plin5 | Integrated DNA Technologies | Mm.PT.56a.8290728 | n/a |
| Ptk2 (FAK) | Integrated DNA Technologies | Mm.PT.58.30114478 | n/a |
| Ppia | Integrated DNA Technologies | Mm.PT.39a.2.gs | n/a |
| Pparg | Integrated DNA Technologies | Mm.PT.58.31161924 | n/a |
| Ppargc1a | Integrated DNA Technologies | Mm.PT.58.16192665 | n/a |
| Antibodies | | | |
| Plin5 | Thermo Fisher Scientific | Cat# PA1-46215 | 1:1000 |
| UCP1 (E9Z2V) XP Rabbit mAb | Cell Signaling Technology | Cat# 72298 | 1:500 |
| b-tubulin | Cell Signaling Technology | Cat# 2146 | 1:1000 |
| b-actin | Cell Signaling Technology | Cat# 4967 | 1:1000 |
| MRTFA | Cell Signaling Technology | Cat# 97109 | 1:1000 |
| phospho-FAK (Tyr397) | Cell Signaling Technology | Cat# 3283 | 1:1000 |
| FAK | Cell Signaling Technology | Cat# 3285 | 1:1000 |
| phospho-CREB (Ser133) (87G3) | Cell Signaling Technology | Cat# 9198 | 1:1000 |

| | | | |
|---|---|---|---|
| CREB | Cell Signaling Technology | Cat# 9197 | 1:1000 |
| phospho-HSL (Ser563) | Cell Signaling Technology | Cat# 4139 | 1:1000 |
| HSL | Cell Signaling Technology | Cat# 4107 | 1:1000 |
| Myh9 (Myosin IIa) | Cell Signaling Technology | Cat# 3403 | 1:1000 |
| Myh10 (Myosin IIa) | Cell Signaling Technology | Cat# 3404 | 1:1000 |
| OPA1 | Abcam | Cat# ab42364 | 1:1000 |
| MFN2 | Cell Signaling Technology | Cat# 9482 | 1:1000 |
| phospho-Drp1 (Ser616) | Cell Signaling Technology | Cat# 3455 | 1:1000 |
| Drp1 | Cell Signaling Technology | Cat# 8570 | 1:1000 |
| IRDye 680LT Goat anti-Mouse IgG antibody | LI-COR Biosciences | Cat# 925-68020 | 1:10000 |
| IRDye 800CW Goat anti-Rabbit IgG antibody | LI-COR Biosciences | Cat# 925-32211 | 1:10000 |
| Imaging Reagents | | | |
| Lipi-Blue | Dojindo | Cat# LD01 | 0.1 mM |
| PKMito deep red | cytoskeleton | Cat# CY-SC055 | 1: 1000 (vendor protocol) |
| SPY555-Fast Act | cytoskeleton | Cat# CY-SC205 | 1: 1000 (vendor protocol) |
| SPY650-FastAct | cytoskeleton | Cat# CY-SC505 | 1: 1000 (vendor protocol) |
| DAPI | Thermo Scientific | Cat# D1306 | 1 mg/ml |
| BODIPY™ 505/515 | Thermo Scientific | Cat# D3921 | 1 mg/ml |
| Fluo-4 AM | Invitrogen | Cat# F14201 | 3 mM |
| Pluronic F-127 | Invitrogen | Cat# P3000MP | 0.10% |
| Public Dataset | | | |
| Cold exposed mouse iWAT | Holman, C. D. et al. | GSE227441 | |

**Supplemental table 1. List of chemicals and reagents.**
